# The Product neutrality function defining genetic interactions emerges from mechanistic models of cell growth

**DOI:** 10.1101/2024.11.29.626097

**Authors:** Lucas Fuentes Valenzuela, Paul Francois, Jan Skotheim

## Abstract

Genetic analyses, which examine the phenotypic effects of mutations both individually and in combination, have been fundamental to our understanding of cellular functions. Such analyses rely on a neutrality function that predicts the expected phenotype for double mutants based on the phenotypes of the two individual non-interacting mutations. In this study, we examine fitness, the most fundamental cellular phenotype, through an analysis of the extensive colony growth rate data for budding yeast. Our results confirm that the Product neutrality function describes the colony growth rate, or fitness, of a double mutant as the product of the fitnesses of the individual single mutants. This Product neutrality function performs better than additive or minimum neutrality functions, supporting its continued use in genetic interaction studies. Furthermore, we explore the mechanistic origins of this neutrality function by analyzing two theoretical models of cell growth. We perform a computational genetic analysis to show that in both models the product neutrality function naturally emerges due to the interdependence of cellular processes that maximize growth rates. Thus, our findings provide mechanistic insight into how the Product neutrality function arises and affirm its utility in predicting genetic interactions affecting cell growth and proliferation.

## 1 Introduction

Genetic analysis has been one of the primary methods scientists use to understand how a cell works. One way this is done is through the analysis of genetic interactions, in which the phenotypic effects of mutations are analyzed both as single mutations and then together as double mutations in the same cell. Genes are then said to interact if their combination produces phenotypes that are different from what is predicted from a generic model combining two non-interacting mutations (Phillips 2008; Beltrao et al. 2010; Costanzo, Kuzmin, et al. 2019). In other words, a genetic interaction is identified when combining multiple mutations yields something unexpected. Yet, what should we expect when combining mutations in such a complex system as a living cell?

The expected phenotype of a double mutant predicted from the two single mutants’ phenotypes is defined by the neutrality function. In this way, the neutrality function calculates the expected phenotype of a double-mutant strain carrying two non-interacting mutations (Mani et al. 2008; Beltrao et al. 2010). If the double mutant phenotype deviates significantly from that given by the neutrality function for two specific mutations, they are then said to interact. A lot of care therefore needs to be taken in selecting the appropriate neutrality function, which depends on the context and phenotype to be examined (Phillips 2008). The neutrality function should be defined such that most mutations are categorized as non-interacting. If the neutrality function were not defined this way, the majority of genes would appear to interact, leaving only a few genes with distinct functions. However, decades of cell biological and structural biological analysis has identified specific functions for many genes and their associated proteins. For example, the components of the ribosome or RNA polymerase have the specific task to form these complexes, and metabolic enzymes catalyze specific biochemical reactions. This implies that a judiciously selected neutrality function should predict most double mutant fitnesses from the single mutant fitnesses since any two randomly selected genes should be unlikely to interact. Recent technological advances have enabled the screening of the proliferation of single and double genetic mutants at increasingly larger scales in *E. coli* (Typas et al. 2008; Butland et al. 2008; Babu et al. 2014), fission yeast (Roguev et al. 2008; S. J. Dixon et al. 2008), C. elegans (Lehner et al. 2006; Byrne et al. 2007), and human cells (Horlbeck et al. 2018). In budding yeast, Synthetic Genetic Arrays (SGA) (Tong et al. 2001) have generated the largest and most comprehensive such datasets (Baryshnikova, Costanzo, Kim, et al. 2010; Costanzo, Baryshnikova, et al. 2010; Costanzo, VanderSluis, et al. 2016).

The most fundamental phenotype of a cell is its fitness, namely how quickly it grows and proliferates in a given environment. Here, we focus on this property and define fitness as the relative exponential growth rate with respect to that of a wild type cell. For yeast proliferation, single mutant fitnesses are usually assumed to combine according to a Product neutrality function, namely that the fitness of the double mutant is the product of the fitnesses of the single mutants (Mani et al. 2008). This neutrality function was shown to better predict double mutant fitnesses than an Additive neutrality function, where the differences between wild type and mutant fitnesses were simply added together, and a Minimum neutrality function, where the double mutant fitness was taken as the lowest fitness of the two single mutant strains. However, this analysis was based on older, less extensive data, which raises the question of whether this neutrality function remains accurate when the large amount of more recently collected yeast data is also considered. And, if so, then why does the Product neutrality function accurately describe double mutant fitnesses? In other words, what are the properties of the underlying system controlling cell growth and proliferation that result in a Product neutrality function for mutant fitnesses?

In this paper, we conduct an in-depth analysis of recent yeast double mutant datasets and show that they support the Product neutrality function. Moreover, we analyze two theoretical models of growth of increasing complexity (Scott, Gunderson, et al. 2010; Weiße et al. 2015) and find that the Product neutrality function emerges naturally from both growth models, albeit with small deviations specific to each. Taken together, our work supports the use of the Product neutrality function to model genetic interactions in the regulation of cell growth and gives mechanistic insight into it origin.

## 2 Results

### 2.1 High-throughput gene perturbation experiments in budding yeast support a Product neutrality function for double mutant fitness

To test the general validity of neutrality functions for mutations affecting cell proliferation, we sought to examine the most extensive such dataset. In the Synthetic Genetic Array (SGA) dataset, over 20 million single and double mutant budding yeast strains were generated. Then, the growth rates of their colonies were measured in synthetic genetic arrays (Tong et al. 2001; Baryshnikova, Costanzo, Kim, et al. 2010; Costanzo, Baryshnikova, et al. 2010; Costanzo, VanderSluis, et al. 2016). Each mutant’s fitness was then defined as this measured growth rate normalized by that of the wild-type strain, enabling a consistent comparison across thousands of genotypes. Next, we use SGA data sets of growth of single and double mutant cells for pairs of mutations to test specific neutrality functions.

Here, we followed (Mani et al. 2008) and began our examination using the Product, Additive, and Minimum neutrality functions (see Fig. 1A). The Product neutrality function predicts that the fitness of a double mutant is the product of the fitnesses of the two corresponding single mutants, while the Additive neutrality function proposes that the difference between the double mutant and wild type fitnesses is the sum of the differences between the two mutant and wild type fitnesses. The Minimum neutrality function proposes that the fitness of the double mutant is equal to the fitness of the least fit single mutant. Effectively, these neutrality functions express different forms of modularity or independence between cellular processes. The Product model suggests that a mutation’s effect depends on the fitness of the background strain without that mutation, while a mutation’s effect is independent of the fitness of the background strain in an Additive model. The Minimum model suggests that there is some rate-limiting process whose slow time scale dominates the determination of cell growth so that more minor mutations affecting other processes have no additional effect.

**Figure 1:**
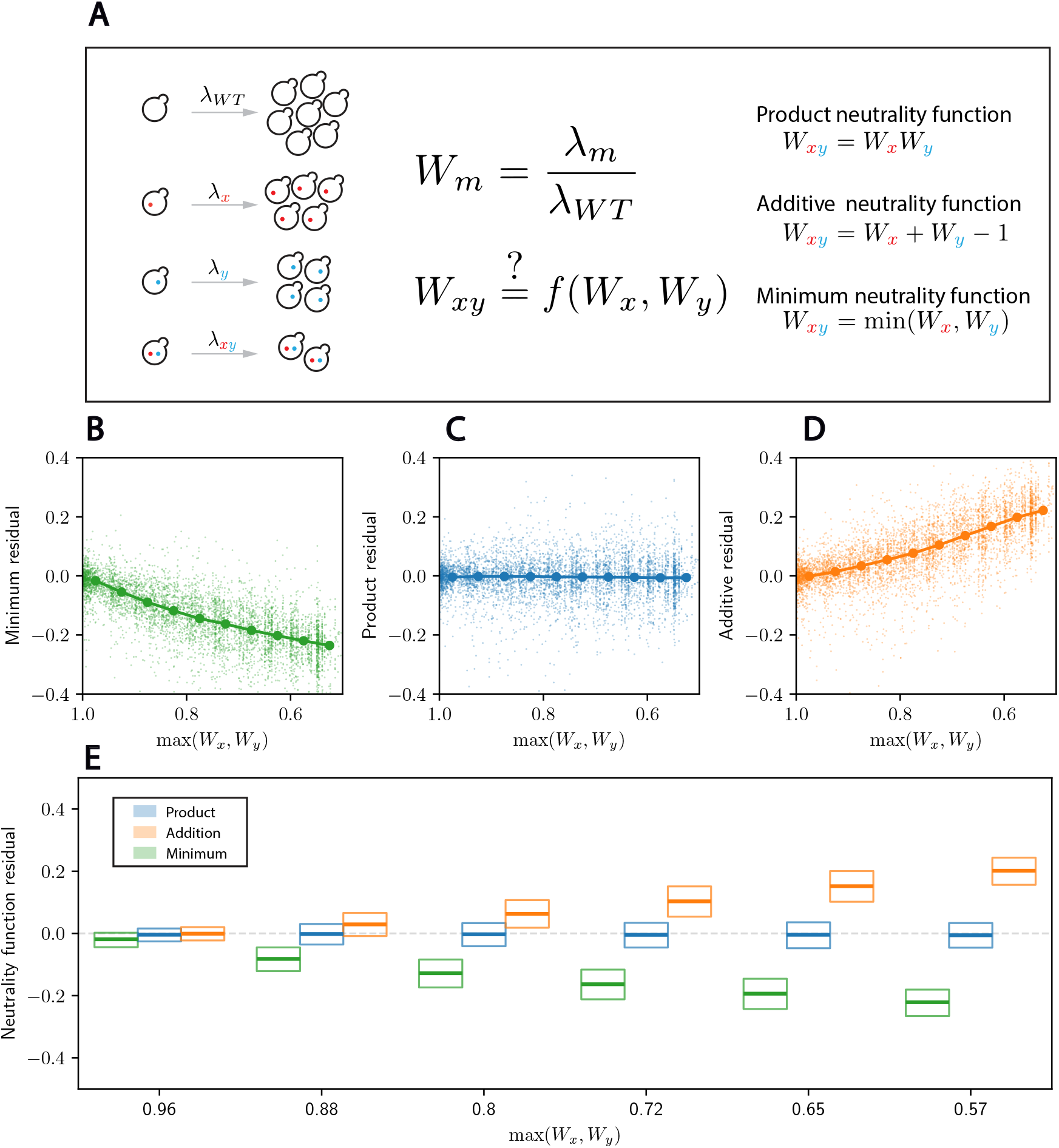
High-throughput gene deletion experiments in budding yeast support a Product neutrality function for double mutant fitness. **A.** Budding yeast mutant fitness is defined as the colony growth rate relative to that of wild type cells. Schematic illustration of epistasis in growth rate, and of different laws proposed in the literature. *λ* denotes the growth rate, and W the fitness. **B-D**. For each double mutant, we plot the residual of the fitness predicted from the indicated model against the fitness of the fittest of the two separate single mutants (maximum single mutant fitness). Dots indicate the median for 10 equally spaced bins between 0.5 and 1. **E**. Box plots for the distributions of the residuals for the three neutrality functions as a function of the maximum single mutant fitness. Thick line denotes the median, and boxes denote the 25th and 75th percentiles of the distributions. The data plotted here represents a subset of the entire SGA dataset, corresponding to the Deletion Mutant Array (DMA) at 30°C. Figs. S1& S2 report results for the other sub-datasets.

To compare the different neutrality functions with double mutant fitnesses, we first perform some minor pre-processing of the SGA data (see Methods for details). We note that these data are from combinations of gene deletions, temperature-sensitive alleles, and hypomorphic mutants (see Methods). Moreover, cells were growing quickly on the relatively rich synthetic complete media containing glucose (Baryshnikova, Costanzo, S. Dixon, et al. 2010). For these reasons, increasing the growth rate is difficult and we are in the regime where mutations generally decrease fitness. This contrasts to evolution experiments, where fitness increases very slowly through the gradual accumulation of mutations, which likely exhibit different neutrality functions from those we consider here (Phillips 2008; Bakerlee et al. 2022; Johnson et al. 2023). Consistent with previous work (Mani et al. 2008), we see that the Product neutrality function better predicts double mutant fitnesses as a function of the single-mutant fitnesses over a broad range of fitness defects (Fig. 1B-E, S1). Indeed, the median residual for the Product neutrality function remains very close to zero even for highly deleterious mutations, while it significantly deviates for the other two. For instance, for a maximum single mutant fitness of 72%, the median residual is −0.8 % for the Product neutrality function, while they are −16.8 % and 9.8% for the Minimum and Additive neutrality functions, respectively. For smaller values of the maximum single mutant fitness, the median residual remains virtually unchanged for the Product neutrality function, while it deviates even further from zero for the other two. However, we note that significant variation around the median residual remains. At a maximum single mutant fitness of 72%, the interquartile range lies between 8% and 10% depending on the neutrality function considered.

This observation holds for mutants carrying mutations to essential and nonessential genes, and across different temperature conditions (see Methods and Figs S1 & S2). The Minimum neutrality function generally predicts fitnesses that are too high, while the Additive neutrality function generally predicts fitnesses that are too low.

### 2.2 The Product neutrality function describes interactions between genes associated with two distinct biological processes

While the Product neutrality function predicts double mutant fitnesses better than the other ones we considered, there remains significant variation (residuals) in the data. This suggests that mutations affecting different functional parts of the cell might be following different neutrality functions. To determine whether this is the case, we analyze the distribution of epistasis residuals for pairs of distinct biological processes and their associated genes. We use the Gene Ontology (GO) dataset (Ashburner et al. 2000; Gene Ontology Consortium et al. 2023) and, for each biological process, extract the genes in the SGA dataset that are associated with this process (see Methods for details). In particular, we define inter-process gene pairs as pairs of gene perturbations in two distinct biological processes identified using GO annotations (Fig. 2A). Similarly, intra-process pairs are defined as pairs of gene perturbations in the same GO-annotated biological process (Fig. S3A). Then, for each pair of GO-defined processes, we compute the residuals for the neutrality functions for all pairs of mutations where one mutation is associated with one process and the second mutation with the other. For each neutrality function and pair of GO processes, we extract the median residual as a function of the largest single mutant fitness defect (a proxy for mutation severity). This shows that, while imperfect, the Product neutrality function is a good description of typical interactions, while the Additive and Minimum neutrality functions have large, systematic residuals (Fig. 2B-D). In general, this result is expected and consistent with the use of this type of genetic analysis to define mutations in genes from different biological processes as not interacting. Moreover, we find that this is not only generally true, but also true for each specific pair of processes that we consider. In other words, we do not find evidence that there are particular pairs of processes whose mutations significantly deviate from the Product neutrality function or more closely follow an Additive or Minimum neutrality function.

**Figure 2:**
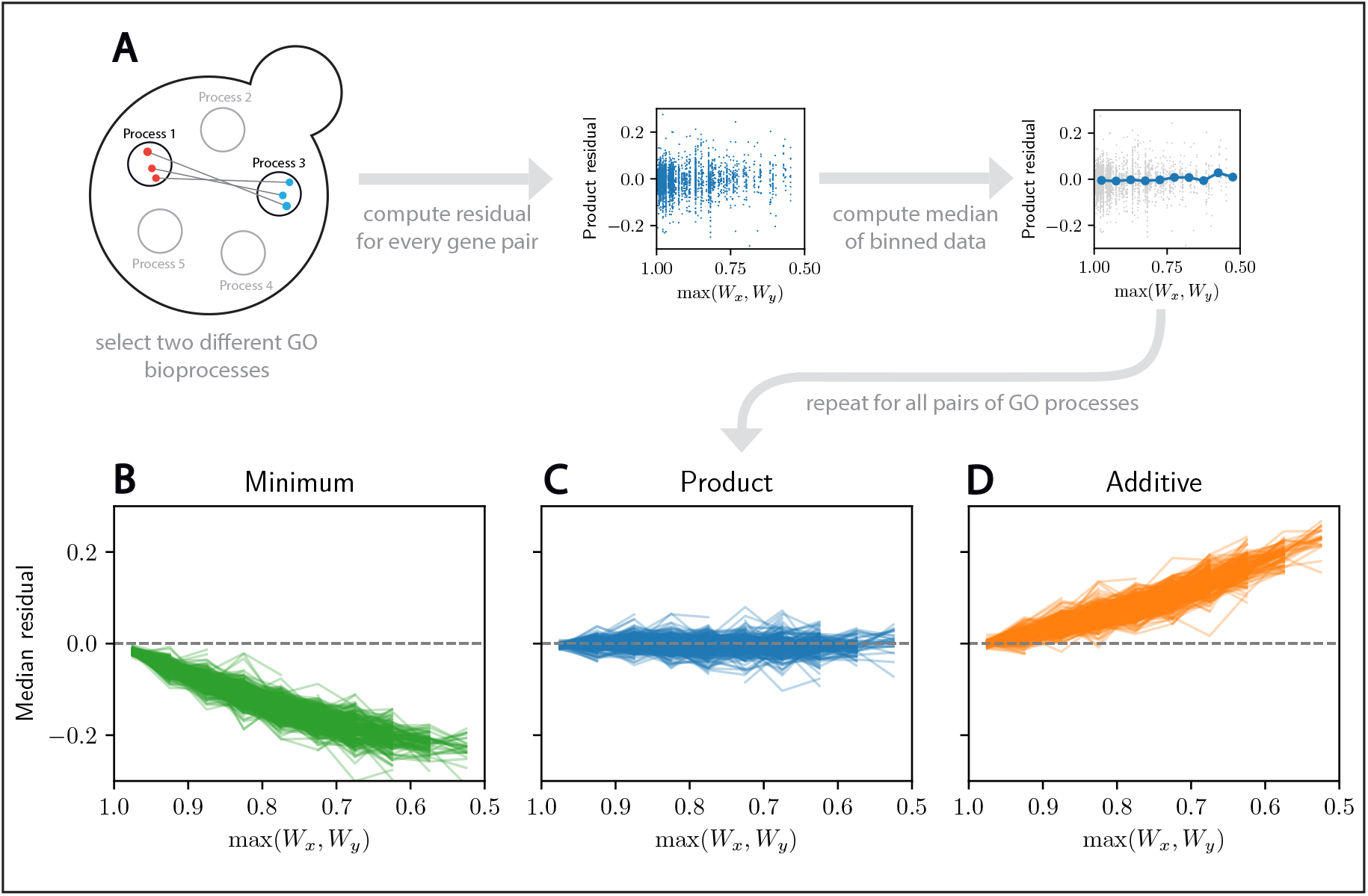
The Product neutrality function describes interactions between genes associated with two distinct biological processes. **A.** Schematic illustration of the analysis process. We first select two different GO biological processes and extract the double mutants in the SGA dataset associated with them. Then, we compute the median residual for each pair of biological processes and each neutrality function. **B-D**. Median residual for the Minimum, Product, and Additive neutrality functions as a function of the maximum single mutant fitness. Each line denotes mutations to a different pair of distinct GO biological processes. The majority of biological process pairs closely follow the Product model.

While mutations associated with different biological processes have fitnesses generally predicted by the Product neutrality function, since they generally do not interact, this may not be the case for mutations associated with the same process. For example, if two mutations break the same protein complex, one would not expect any additional drop in fitness for the double mutant. Consistent with this notion, for gene pairs in the same biological process, we observe more deviations as well as significantly larger residuals (Fig. S3B-D). The quantitative comparison of the two types of interactions reveals that large residuals (both positive and negative) are significantly more likely for two mutations categorized as being in the same GO process (Fig. S3E-F).

We note that the SGA dataset has already been used to assign a biological process to each gene (Costanzo, VanderSluis, et al. 2016). For each mutation, a vector of residuals for the Product neutrality function with all other mutations was generated. Then, a Pearson correlation coefficient was calculated for each pair of these vectors. The reasoning was that mutations affecting the same biological process should have similar genetic interaction profiles, which was found to be the case. This then allowed the clustering of groups of correlated mutations, which were named using prior knowledge of many genes in each cluster. That this analysis generally works, i.e., the gene clusters have discernible biological meaning, can be viewed as further support of the Product neutrality function.

### 2.3 A bacterial growth model partially supports the Product neutrality function

Having verified empirically that the Product neutrality function is supported by the latest data for cell proliferation, we now turn our attention to its origins. Addressing this question requires some mechanistic model of biosynthesis. However, most mechanistic models of growth apply directly to single cells in rich nutrient conditions, which may not directly apply to the SGA measurements of colony expansion rates. In particular, colony growth has been shown to follow a biphasic pattern (Meunier et al. 1999). A first exponential phase is followed by a slower linear phase as the colony expands. Previous modeling and empirical work indicates that this second linear expansion rate reflects the underlying exponential growth of cells in the periphery of the colony (Pirt 1967; Gray et al. 1974; Gandhi et al. 2016; Baryshnikova, Costanzo, S. Dixon, et al. 2010; Zackrisson et al. 2016; Miller et al. 2022). More precisely, mathematical models show the linear colony-size expansion rate is directly proportional to the square root of the exponential growth rate under non-limiting conditions. Intuitively, this relationship arises because colony growth is dominated by the expansion of the population of cells in an annulus at the colony border that are exposed to rich nutrient conditions. These cells expand at a rate similar to the exponential rate of cells growing in a rich nutrient liquid culture. In contrast, the cells in the interior of the colony experience poor nutrient conditions, grow very slowly, and do not contribute to colony growth.

This intimate relationship between both proliferation rates allows us to explore the origin of the Product neutrality function in mechanistic models of cell growth. Indeed, if colony-based fitnesses follow a Product model, then

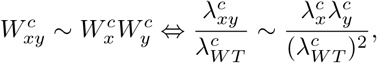

where the superscript *c* indicates colony-based values for the fitness *W* and the growth rate *λ*. Taking into account the relationship between single-cell exponential growth rates and colony growth rates, we can write

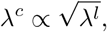

where the superscript *l* denotes liquid cultures. Combining these expressions, we obtain

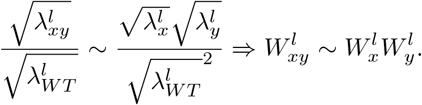

In other words, from the perspective of the Product neutrality function, fitnesses based on colony expansion rates are equivalent to fitnesses based on single-cell exponential growth rates. The prevalence of the Product neutrality model—both in the SGA data and in previous studies on datasets from liquid cultures (Jasnos et al. 2007; Onge et al. 2007; Mani et al. 2008)—encourages the exploration of its origin in mechanistic models of cell growth.

While models of entire cells do exist, these models are complex and computationally intensive (Karr et al. 2012; Oftadeh et al. 2021). This makes probing and extracting explanatory information from these models difficult. We therefore sought to analyze simpler, more tractable, lower-dimensional models of cell growth. Coarse-grained models offer an appealing alternative for probing the fundamental principles of metabolism and growth (Scott, Gunderson, et al. 2010; Roy et al. 2021; Weiße et al. 2015; Balakrishnan et al. 2022; Chure et al. 2023; Calabrese et al. 2023). Rather than representing as many reactions as possible, they provide an integrated representation of generic processes in the cell. Their simplicity and low-dimensionality makes them easy to compare with empirical measurements and to examine for potential explanatory relationships.

The reduced, tractable model of cell growth that we will consider first was developed for *E. coli* (Scott, Gunderson, et al. 2010; Scott and Hwa 2011) (Fig. 3A). While this model was developed for *E. coli* bacteria, and validated using data from this organism, there is nothing specific to prokaryotes in the model. Experimental measurements in other organisms suggest that the observations leading to this model, including that the cellular ribosome fraction increases with growth rate, are in fact generic and also seen in the yeast *S. cerevisiae* (Metzl-Raz et al. 2017; Elsemman et al. 2022; Xia et al. 2022). In its simplest form, the model defines growth as resulting from two sets of processes, metabolic and translational, that interact in a linear pathway. The metabolic sector provides precursors that are then assembled into proteins by the translational sector and the flux through each sector is determined by the amount of proteins in that sector. For optimal growth, in which no proteins are wasted, the flux through the metabolic sector is equal to the flux through the translational sector so that

**Figure 3:**
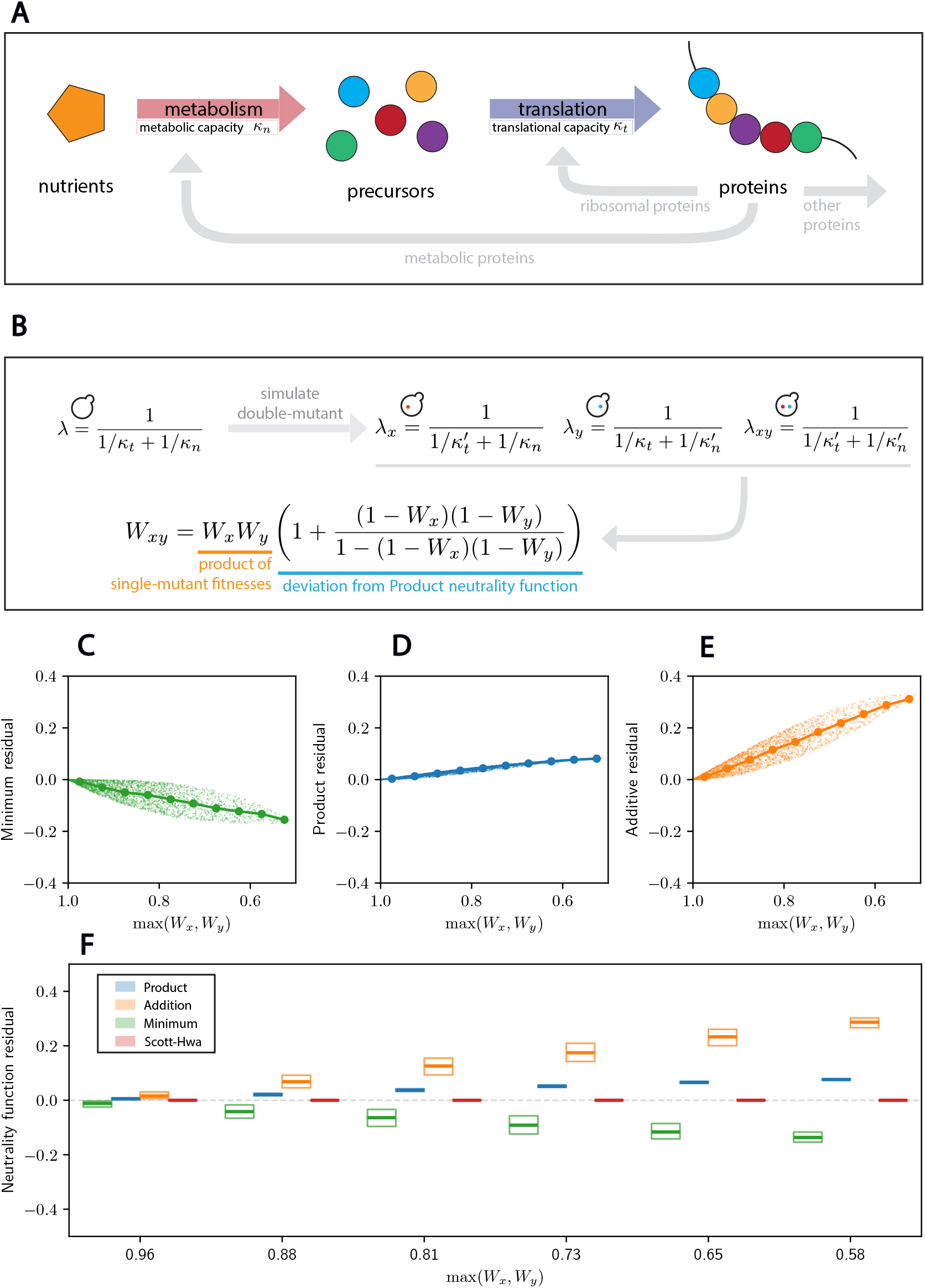
A bacterial growth model partially supports the Product neutrality function. **A**. Schematic of the bacterial growth model by Scott and Hwa. Growth rate is defined by the translation flux, which is itself equal to the metabolic flux. The cell partitions its proteome so as to maximize growth rate. **B**. Mutations are modeled such that they affect either of the parameters, separately. Values of *κ*_*t*_ and *κ*_*n*_ in the mutant are indicated with primes and are sampled from a uniform distribution from 0 to their value in wild type cells. *λ* indicates the corresponding growth rate. The analytical expression of the double-mutant fitness consists of the product model with a perturbation. **C-E**. For each sampled double-mutant, we plot the residual of the fitness predicted from the indicated model against the fitness of the fittest of the two separate single mutants (maximum single mutant fitness). Dots indicate the median for 10 equally spaced bins between 0.5 and 1. **F**. Box plots for the distributions of the residuals for the three models and the model in C as a function of the maximum single mutant fitness. Thick line denotes the median, and boxes denote the upper and lower quartiles of the data. The analytical model in B, named Scott-Hwa and shown in red, is exact.

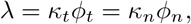

where the flux through translation and metabolic sectors are characterized by the parameters *κ*_*t*_ and *κ*_*n*_ multiplying the fraction of the proteome devoted to ribosomes and metabolic proteins, *ϕ*_*t*_ and *ϕ*_*n*_ respectively. These fluxes directly determine the growth rate of the cell, *λ*. The total proteome is fixed and is partitioned into metabolic, translational, and “other” sectors of the cell so that

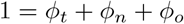

where *ϕ*_*o*_ is the fraction of the cell devoted to other housekeeping functions. This set of algebraic equations can be solved for the optimal growth rate

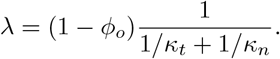

We can then model a mutation as a perturbation to these parameters that decreases the growth rate since these are the types of mutations that dominate the budding yeast data (Fig. 3B). Single mutants have either *κ*_*t*_ or *κ*_*n*_ perturbed, while double mutants have both parameters perturbed. Given that we are concerned with two non-interacting mutations, we do not consider the cases where both mutations affect the same parameters. Indeed, one expects two mutations affecting the same parameter to interact. These combinations are therefore inappropriate to study the emerging neutrality functions from growth models, and we do not consider them in this paper.

We can also consider mutations to *ϕ*_*o*_, which could be associated with deleting a gene encoding a protein that is not required for the given growth condition. This would serve to increase the cell growth rate because now a larger fraction of the proteome could be devoted to metabolism and translation. In this case, a mutation to *ϕ*_*o*_ and another to either *κ*_*t*_ or *κ*_*n*_ would combine exactly multiplicatively so that

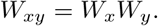

However, in the generally rich media conditions the SGA experiments were done, there is no evidence that any gene deletion causes an increase in cell growth rate so we do not consider this type of mutation further. We therefore ignore the multiplicative factor (1 ™ *ϕ*_*o*_) and analyze the following expression for growth rate

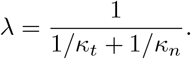

We can then analytically derive a closed-form solution for the double-mutant fitness as a function of the single-mutant fitnesses (Fig. 3C; see SI for details). Under this model, which we call Scott-Hwa in reference to the authors of the initial work, we observe that the Product neutrality function fits the mutational analysis of the Scott-Hwa model better than the Additive or Minimum neutrality functions (Fig. 3D-G).

We understand the better performance of the Product neutrality function to arise from a type of feedback regulation that ensures that the flux through all sectors be equal. This effectively makes the mutations to the metabolic and translational sector interdependent, despite their *a priori* independent functions. In this model, these sectors are coupled because the cell is assumed to have a feedback process to optimize growth rate under any perturbation to its parameters. For instance, in the case of a mutation that decreases the metabolic capacity *κ*_*n*_, this feedback drives an increase in the fraction of metabolic proteins at the expense of translational proteins such that the growth rate is maximized under those new parameters. If this feedback were absent, then the growth rates of the double mutant would be significantly lower. In that case, the double-mutant fitness actually follows a Minimum neutrality function (see supplementary information and Fig S4). Finally, we also note that the Product neutrality function does not accurately predict model fitnesses well for beneficial mutations—i.e. mutations that increase growth rate (see Fig. S5). This is because the deviations from the Product neutrality function in the mathematical derivation for the double-mutant fitness in Fig. 3 can diverge for beneficial mutations (e.g. *W*_*x*_ = *W*_*y*_ = 2). When the Scott-Hwa model’s growth-optimizing feedback operates in the context of beneficial mutations, as one process is made more efficient, proteomic resources are allocated to accelerate other processes in the cell. In this way, improving the efficiency of one process will indirectly benefit other processes, leading to compound effects such that double-mutant fitnesses are higher than any of the three models predicts for beneficial mutations.

We note that in this analysis we do not aim to replicate the statistics of mutations in the SGA dataset, where mutations to either sector could be statistically rarer or more frequent than the other. Instead, we here aim to analyze how mutations to independent parameters governing cell growth combine considering the simplest model with two sectors and their corresponding parameters.

### 2.4 The Product neutrality function accurately predicts fitness for many pairs of parameters in a more complex cell growth model

While the Scott-Hwa model has proven successful for predicting many aspects of bacterial growth, it remains very simple. Therefore, we sought to explore a more complex model that explicitly incorporates more aspects of biosynthesis. Here, we consider the model of (Weiße et al. 2015), which incorporates nutrient intake, transcription, competitive binding between mRNAs and ribosomes, and translation, all of which are mediated by associated enzymes and a limiting cellular “energy” (Fig. 4A). The Weiße model decomposes cell growth into multiple steps (see supporting material for a full model description). External nutrients are first imported into the cell and then metabolized into a cellular “energy”. Both of these steps are catalyzed by associated transport and metabolic enzymes according to Michaelis-Menten kinetics. Transcription and translation are then activated by this generated “energy”, also via Michaelis-Menten kinetics. In particular, the model incorporates different transcription rates for ribosomal and non-ribosomal mRNAs. Different mRNAs then compete for free ribosomes to form a ribosome-mRNA complex. This mRNA competition is modeled using mass action kinetics with specified binding and unbinding rates. Four types of proteins are explicitly modeled as a product of translation: transport proteins, metabolic enzymes, ribosomal proteins, and so-called q-proteins which support housekeeping functions much like the “other” proteins in the Scott-Hwa model.

**Figure 4:**
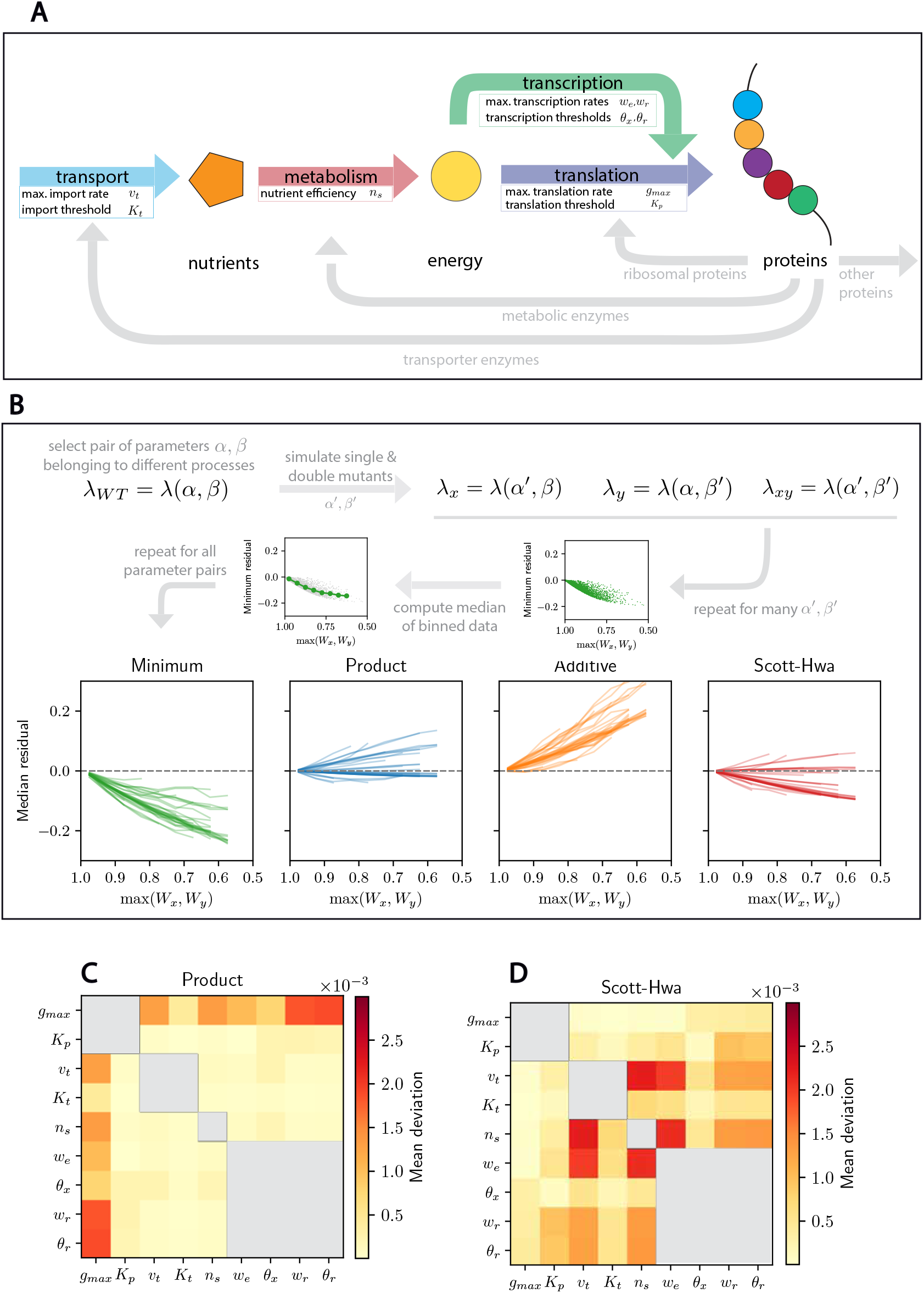
The Product neutrality function accurately predicts fitness for many pairs of parameters in a more complex cell growth model. **A.** Schematic of the growth model from Weiße et al. 2015. This model includes nutrient intake, metabolism, transcription, and translation. **B**. Schematic of the mutational analysis. For each pair of parameters *α* and *β*, mutations are modeled such that they affect either of the parameters, separately. Then, the median residual is computed for each neutrality function and they are subsequently reported for every pair of parameters considered. **C-D**. For each parameter pair, we report the mean deviation of the simulated double mutants from (C) the Product neutrality function and (D) the analytical expression of the double mutant fitness under the Scott-Hwa model. Only parameter pairs corresponding to two different biological processes are considered. Those corresponding to the same process are greyed out. Parameter pairs involving translation (*g*_*max*_, *K*_*p*_) are the ones described best by the Scott-Hwa model while the others are better described by the Product neutrality function.

To perform a mutational analysis of the Weiße model, we first identified the parameters in the model that can reasonably be expected to change through a gene perturbation (see supporting material for details). For instance, we assume that some parameters, such as the maximum nutrient import rate, could be impacted by a gene perturbation, while other parameters, such as the average gene length in the genome, could not. This led to the identification of 9 easily interpreted parameters whose mutation could negatively impact the cell growth rate and corresponding to 28 parameter pairs associated to different biological processes. We then performed a similar analysis as we did for the Scott-Hwa model. Namely, we constructed mutants for each pair of parameters by rescaling the values of the original parameters by a number randomly sampled between 0 and 1 (see Methods for details). We then analyzed the statistics of the epistasis coefficients for the neutrality functions that we considered so far (Fig. 4B).

Our mutational analysis of the Weiße model revealed several striking observations. First, the Product neutrality function is generally better than the Additive or Minimal neutrality functions at describing the mutational results. However, we observe a range of responses, and can identify two key subpopulations of parameter pairs. The subset of parameter pairs involving protein translation follow the Scott-Hwa model very closely (Fig. 4D). This is not entirely surprising, as the Weiße model is an extension of the Scott-Hwa model and incorporates a similar global feedback optimizing cell growth and a competition for resources. On the other hand, another subset of parameter pairs follows the Product neutrality function even more closely. These parameter pairs involve mutations to the other sectors, including the transport, metabolism and transcription sectors (Fig. 4C). This raises the question of why some parameter pairs more closely follow the Product neutrality function than others.

### 2.5 Nonlinear kinetics drive deviations from the Product neutrality function in the Weiße model for cell growth

To address the question as to what drives deviations from the Product neutrality function in our genetic analysis of the Weiße model, we sought to take an analytical approach. We examined the dependence of the growth rate on the parameter pairs exhibiting small deviations from the Product neutrality function. To do this, we first extract a closed form expression that models the growth rate *λ* and its dependence on two mutated parameters *α* and *β* (Fig. 5A; see supporting material). While only approximate, this derivation represents the data appropriately for members of this subset of parameters (see Fig. S7). There are two striking features in this derivation. First, the growth rate *λ* has an explicit dependence on the two parameters *α* and *β*, when *α* and *β* are selected from the subset of parameters governing metabolism and transport sectors. Second, the amplitude of deviation from the Product neutrality function is governed by an additional parameter, *γ*, which is the inverse of the Michaelis-Menten constant giving the transcription rate as a function of the cellular “energy”. Thus, when *γ* is small, transcription is a less efficient process that is then linearly related to the cellular “energy” available. When *γ* is larger, transcription is saturated and performed at a rate unrelated to the available “energy”. As we decrease *γ*, we observe that the Product neutrality function is a better and better approximation (Fig. 5C). Importantly, the intuition provided by the analytical approximation extends to multiple pairs of parameters (see Supplementary Information and Fig. S8). Taken together, our analysis of the Weiße model shows how the Product neutrality function naturally arises for many different parameter pairs and how deviations from it can be driven by nonlinear effects, such as those that can emerge from Michaelis-Menten kinetics.

**Figure 5:**
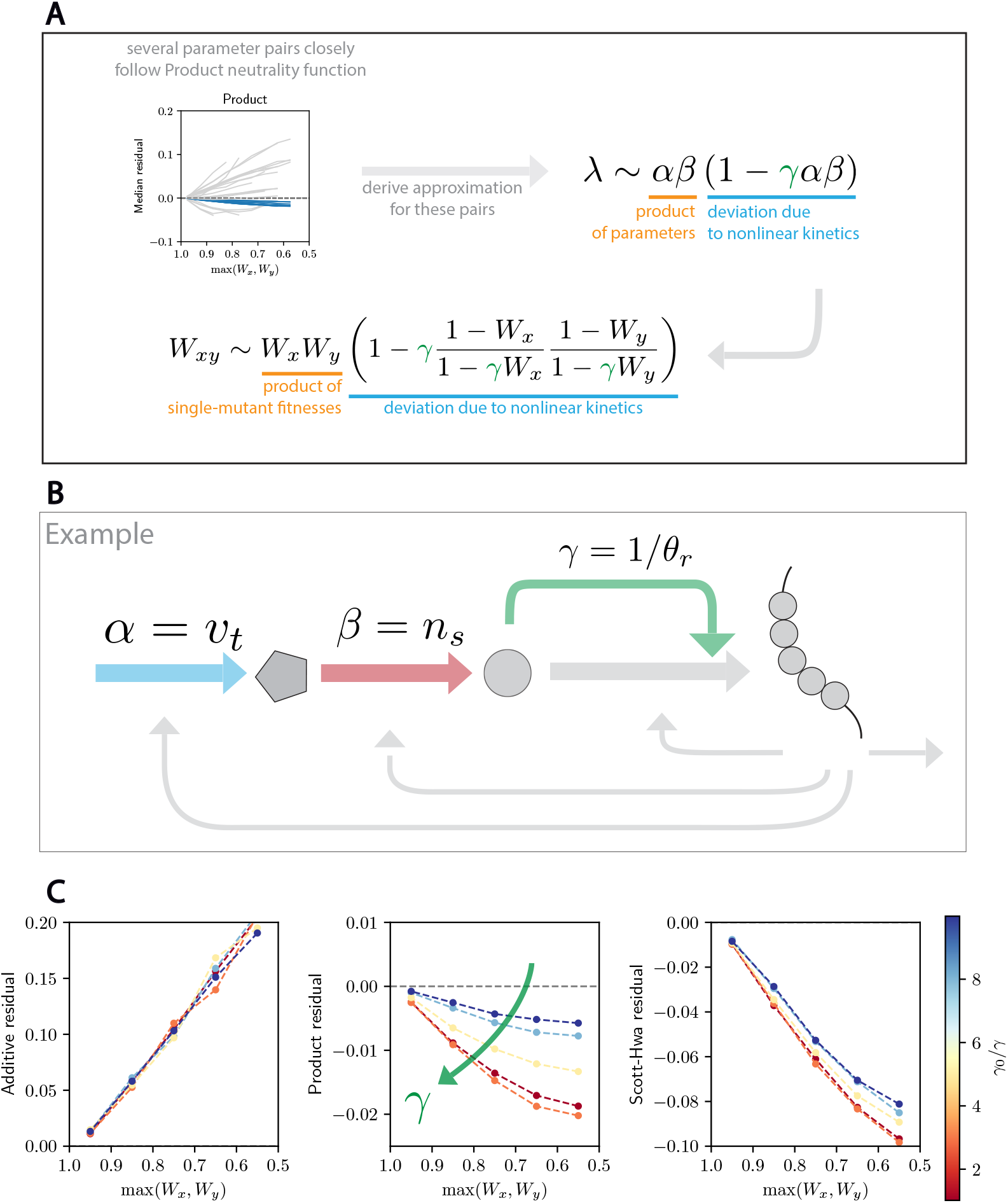
Nonlinear kinetics drive deviations from the Product neutrality function in the Weiße model. **A.** A subset of parameter pairs we analyzed follow the Product model very closely. We derived an analytical approximation of the growth rate and the double mutant fitness for these pairs, and found that the deviation from the product law is governed by nonlinear kinetics. **B**. In the case of the parameter pair (*v*_*t*_, *n*_*s*_) we show that the deviation from the Product model is driven by the Michaelis-Menten constant *θ*_*x*_ associated with transcription (see Supporting Information). **C**. Tuning the value of *γ* impacts how good of an approximation the Product neutrality function is for this and other parameter pairs (see text). This analysis validates the analytical approximation, and highlights how nonlinear kinetics, in this case Michaelis-Menten kinetics, can drive deviations from the Product neutrality function.

## 3 Discussion

Cell growth and proliferation are fundamental to cell biology and have been subject to extensive genetic analysis aiming to understand the underlying regulatory network. Such genetic analysis often aims to identify interactions between mutations through the combination of individual mutations in a double mutant cell. If the double mutant proliferates at an unexpected rate, the mutated genes are considered to interact. This, of course, raises the question as to what is the expected rate of proliferation for a cell containing both mutations given the proliferation rate of a cell containing only one of the individual mutations. By analyzing a high-throughput dataset of interactions of gene perturbations in budding yeast (Costanzo, Baryshnikova, et al. 2010; Costanzo, VanderSluis, et al. 2016), we found that single-mutant fitnesses tend to combine multiplicatively, consistent with earlier work (Mani et al. 2008).

After establishing that the fitness of a double mutant is expected to be approximately the product of the fitness of the individual mutants, namely, the Product neutrality function, we sought to determine if this was also a feature of models of cell growth. If so, then what underlying mechanisms present in these models give rise to this Product neutrality function? Our analysis complements previous, more abstract theoretical attempts at understanding the origin of the Product neutrality function that are not based on any specific model of cell growth (Chiu et al. 2012). Indeed, we found that the Product neutrality function best fits the simulated double-mutant fitnesses despite deviations that depend on the specific parameter pairs and the particular model considered.

That the Product neutrality function fits the budding yeast data and cell growth models better than the Additive and Minimum neutrality functions has important implications for the underlying network controlling cell growth and proliferation. On the one hand, the Minimum neutrality function implies that the double mutant fitness is set by the most deleterious mutation so that the process this gene is involved in becomes rate limiting for cell growth. Clearly, cell growth does not operate this way, likely because the underlying processes are interconnected. Mutations impairing protein translation impact synthesis of all the proteins in the cell so that other processes, like transcription or surface transport, are also affected. As the cell readjusts its machinery to ensure optimal use of resources, interconnected processes are impacted through a redistribution of cellular resources. On the other hand, the Additive neutrality function implies that a mutation has the same absolute effect on the proliferation rate regardless of the presence of another mutation. This is also clearly not the case as the Additive neutrality function consistently predicts fitnesses below those observed in the data. In the models this is due in part to growth-supporting feedback that reapportions the proteome. In reality, this may reflect the presence of the general stress response which supports cells in response to genetic or environmental perturbations limiting their growth rate (Gasch et al. 2000). In this way, the Product neutrality function is a reasonable intermediate model between Minimum and Additive that incorporates — albeit approximately — effects such as growth-optimizing feedback, and is consistent with the phenomenon of *diminishing returns* or *increasing cost* epistasis (Reddy et al. 2021). Moreover, our theoretical analysis gives insight into the mechanistic underpinning of the Product neutrality function. In our analyses, a product of the single mutant growth rates naturally emerges in the analytical treatment of both theoretical models that we consider, albeit with deviation terms that depend on the specific model and simplifying assumptions.

Taken together, our work here constitutes a first step towards understanding the structure of interactions inherent in cell growth models. While we focused on coarse-grained models for their simplicity and mechanistic interpretability, they might be too simple to effectively model large double-mutant datasets and the resulting double-mutant fitness distributions. For instance, it is not possible to differentiate between multiple types of growth rate perturbations impacting the same sector, as they would all be modeled through a limited number of parameters (Metzl-Raz et al. 2017). We therefore expect the combination of high throughput genetic data with the analysis of larger-scale models, for instance based on Flux Balance Analysis, Metabolic Control Analysis, or whole-cell modeling, to lead to important complementary insights regarding the regulation of cell growth and proliferation (Segré et al. 2005; He et al. 2010; Orth et al. 2010; Kacser et al. 1973; Szathmáry 1993; Dykhuizen et al. 1987; Vienne et al. 2023; Karr et al. 2012; Oftadeh et al. 2021; Kryazhimskiy 2021). We also believe that theoretical exploration of fitness landscapes will shed light on the underlying structure of growth and metabolism networks (Guo et al. 2019; Reddy et al. 2021; Boffi et al. 2023). In addition to larger scale models, we see the refinement of the measurement of cell growth rates as a path forward to a better understanding of its regulation (MacLean 2010). While we showed here that the Product neutrality function fits the data well for deleterious mutations, we anticipate that there are significant and meaningful deviations that are currently obscured by experimental noise. Similarly, large-scale measurements of the impacts of beneficial mutations will be instrumental in testing the validity of the Product neutrality function in this other regime. From our modeling efforts, we anticipate that such measurements could give important insights into the underlying genetic network regulating growth and proliferation.

## 4 Methods

### 4.1 Analysis of the SGA dataset

The complete SGA dataset was accessed on [the cell map webpage](https://thecellmap.org/costanzo2016/). In the SGA genetic interaction dataset, a set of query mutant strains is crossed to an ordered array of mutants.

There are two sets of query mutants. The first one consists of a mix of nonessential deletion mutant strains and of temperature-sensitive alleles of essential genes. The second one is a set of mutants carrying hypomorphic, Decreased Abundance by mRNA Perturbation (DAmP) alleles of essential genes.

There are also two types of arrays. The Deletion Mutant Array (DMA) denotes deletions to a set of nonessential genes, while the Temperature Sensitive Array (TSA) contains a mix of essential and nonessential genes.

Both sets of query mutants are crossed to either type of array, at two different temperature conditions, namely 26°C and 30°C. The analysis of Fig. 1 reports the analysis of the first set of query mutants crossed to the DMA at 30°C. In Fig. S1, we report the same analysis for the first set of query mutants for the other array-temperature combinations. In Fig. S2, we report the analysis for the DAmP set of query mutants in the different array-temperature combinations.

For each subdataset — i.e., each combination of query mutants, array, and temperature condition— the data is processed in the following steps. First, only deleterious mutations are kept. That is, we remove mutants having a fitness larger than 1. Second, we eliminate mutants where the additive model predicts a negative fitness, i.e., such that

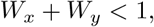

because in this case the prediction under the Additive neutrality function is negative (*W*_*x*_+*W*_*y*_ *−* 1 *<* 0). While we could have analyzed these datapoints with the other neutrality functions, we sought to analyze all neutrality functions on the same consistent dataset.

For the sake of clarity, the scatter plots in Fig. 1 do not reproduce the entire dataset. Instead, the dataset is binned in 10 bins along the x-axis and 500 values are sampled at random in that bin. However, the median lines on top of the scatter plots (e.g. Fig. 1B-D) as well as the box plots (e.g. Figs. 1, S1 and S2) apply to the entire dataset.

### 4.2 Analysis of the GO biological processes

The analysis of GO biological processes is based on the [Uniprot database](https://www.uniprot.org/). In this dataset, genes are associated with a series of GO biological processes. To analyze the behavior of the fitness of double mutants associated with different biological processes, we first selected the set of biological processes that were represented by a large enough number of single mutants in the SGA dataset. Arbitrarily, this limit was set at 50. This led to a limited number of 47 biological processes (and a maximum total of 1081 pairs), which we report in Section 1.1. Naturally, as genes are potentially associated with multiple GO biological processes, this analysis sometimes leads to pairs of processes with genes in common. In this case, we discarded the pair so that we only consider biological process pairs that do not have any genes in common. This results in 685 pairs of biological processes that do not share any genes.

### 4.3 Mutational analysis of growth models

A mutation is modeled as a perturbation of a parameter that decreases the growth rate. For a given parameter *α*, we model a perturbation as *α*^*′*^ = *θα*, where *θ* is a random variable uniformly distributed in [0, 1]. When estimating the impact of the parameter *γ* in the Weiße model (see Section 2.5), we perform a mutational analysis as described above for different values of the parameter *γ*, and collect the median residual.

## 5 Acknowledgements

The authors thank the members of the Mani lab (Northwestern) of the Cremer lab (Stanford University) for insightful discussions and their help in revising the manuscript.

