## supplementary information and figures for "The Product neutrality function defining genetic interactions emerges from mechanistic models of cell growth"

#### 1 Supplementary Information

##### 1.1 Analysis of GO biological processes

The 47 biological processes that result from the analysis (with their GO code) are given in Table S1.

| GO biological process name | GO biological identifier |
|---|---|
| DNA integration | GO:0015074 |
| DNA recombination | GO:0006310 |
| DNA repair | GO:0006281 |
| Ascospore formation | GO:0030437 |
| Cell cycle | GO:0007049 |
| Cell division | GO:0051301 |
| Cell wall organization | GO:0071555 |
| Cellular response to DNA damage stimulus | GO:0006974 |
| Cellular response to oxidative stress | GO:0034599 |
| Chromatin remodeling | GO:0006338 |
| Chromatin silencing at telomere | GO:0006348 |
| Chromosome segregation | GO:0007059 |
| Cytoplasmic translation | GO:0002181 |
| Endocytosis | GO:0006897 |
| Endoplasmic reticulum to Golgi vesicle-mediated transport | GO:0006888 |
| Fungal-type cell wall organization | GO:0031505 |
| Intracellular protein transport | GO:0006886 |
| Intracellular signal transduction | GO:0035556 |
| mRNA splicing, via spliceosome | GO:0000398 |
| Macroautophagy | GO:0016236 |
| Maturation of SSU-rRNA from tricistronic rRNA transcript | GO:0000462 |
| Meiotic cell cycle | GO:0051321 |
| Mitochondrial translation | GO:0032543 |
| Negative regulation of transcription by RNA polymerase II | GO:0000122 |
| Positive regulation of transcription by RNA polymerase II | GO:0045944 |

| | |
|---|---|
| Proteasome-mediated ubiquitin-dependent protein catabolic process | GO:0043161 |
| Protein folding | GO:0006457 |
| Protein import into nucleus | GO:0006606 |
| Protein phosphorylation | GO:0006468 |
| Protein targeting to vacuole | GO:0006623 |
| Protein transport | GO:0015031 |
| Protein ubiquitination | GO:0016567 |
| Pseudohyphal growth | GO:0007124 |
| rRNA methylation | GO:0031167 |
| rRNA processing | GO:0006364 |
| Reciprocal meiotic recombination | GO:0007131 |
| Regulation of transcription by RNA polymerase II | GO:0006357 |
| Regulation of transcription, DNA-templated | GO:0006355 |
| Ribosomal large subunit biogenesis | GO:0042273 |
| Sporulation resulting in formation of a cellular spore | GO:0030435 |
| Transcription by RNA polymerase II | GO:0006366 |
| Transcription elongation from RNA polymerase II promoter | GO:0006368 |
| Translational termination | GO:0006415 |
| Transmembrane transport | GO:0055085 |
| Transposition, RNA-mediated | GO:0032197 |
| Ubiquitin-dependent protein catabolic process | GO:0006511 |
| Vesicle-mediated transport | GO:0016192 |

Table S1: GO biological processes used in the analysis.

#### 1.2 Scott-Hwa model

##### 1.2.1 Derivation of the double-mutant fitness

Here, we derive the expression for the double mutant fitness in the Scott-Hwa model. First of all, the wild-type growth rate $\lambda_{WT}$ is written as

$$\lambda_{WT} = \frac{\kappa_t \kappa_n}{\kappa_t + \kappa_n}. \tag{1}$$

As we consider only deleterious mutations, we can model a perturbation to parameters as

$$\kappa_n' = (1-\delta)\kappa_n, \tag{2}$$
$$\kappa_t' = (1-\varepsilon)\kappa_t, \tag{3}$$

where $\delta, \varepsilon \in [0,1]$. For simplicity, we will denote by the $x$ subscript mutations affecting $\kappa_t$ and the $y$ subscript mutations affecting $\kappa_n$. We note the order does not matter. Therefore, the different fitnesses $W_\cdot = \lambda_\cdot/\lambda_{WT}$ are given by

$$W_x = \frac{1-\varepsilon}{1-\varepsilon\kappa_t/(\kappa_t+\kappa_n)}, \tag{4}$$
$$W_y = \frac{1-\delta}{1-\delta\kappa_n/(\kappa_t+\kappa_n)}, \tag{5}$$
$$W_{xy} = \frac{(1-\varepsilon)(1-\delta)(\kappa_t+\kappa_n)}{(1-\varepsilon)\kappa_t+(1-\delta)\kappa_n}. \tag{6}$$

From there, we can calculate that

$$W_x W_y = \frac{(1-\varepsilon)(1-\delta)(\kappa_t+\kappa_n)}{(1-\varepsilon)\kappa_t+(1-\delta)\kappa_n+\varepsilon\delta\lambda_{WT}}. \tag{7}$$

This expression is similar to Eq. (6). Indeed, we can rewrite it as

$$W_{xy} = W_x W_y \left(1+\frac{\varepsilon\delta\lambda_{WT}}{(1-\varepsilon)\kappa_t+(1-\delta)\kappa_n}\right). \tag{8}$$

We see that the double mutant fitness $W_{xy}$ consists of the product of single mutant fitnesses and a deviation. We will now express this deviation as a function of single mutant fitnesses only, showing that the expression does not depend on the value of the parameters $\kappa_t$ and $\kappa_n$.

Rearranging Eqs (4) and (5), we have

$$1-W_x = \frac{\varepsilon\kappa_n}{(1-\varepsilon)\kappa_t+\kappa_n}, \tag{9}$$

$$1-W_y = \frac{\delta\kappa_t}{(1-\delta)\kappa_n+\kappa_t}, \tag{10}$$

$$\Rightarrow (1-W_x)(1-W_y) = \frac{1}{\kappa_t+\kappa_n}\frac{\varepsilon\delta\lambda_{WT}}{(1-\varepsilon\kappa_t/(\kappa_t+\kappa_n))(1-\delta\kappa_n/(\kappa_t+\kappa_n))}. \tag{11}$$

which can be rearranged to see that

$$\frac{\varepsilon\delta\lambda_{WT}}{(1-\varepsilon)\kappa_t+(1-\delta)\kappa_n} = (1-W_x)(1-W_y)\frac{(1-\varepsilon\kappa_t/(\kappa_t+\kappa_n))(1-\delta\kappa_n/(\kappa_t+\kappa_n))}{(1-\varepsilon)\kappa_t+(1-\delta)\kappa_n} \tag{12}$$

$$= (1-W_x)(1-W_y)\frac{1-\varepsilon\kappa_t/(\kappa_t+\kappa_n)-\delta\kappa_n/(\kappa_t+\kappa_n)+\varepsilon\delta\lambda_{WT}/(\kappa_t+\kappa_n)}{(1-\varepsilon)\kappa_t+(1-\delta)\kappa_n} \tag{13}$$

$$= (1-W_x)(1-W_y)\frac{(1-\varepsilon)\kappa_t+(1-\delta)\kappa_n+\varepsilon\delta\lambda_{WT}}{(1-\varepsilon)\kappa_t+(1-\delta)\kappa_n} \tag{14}$$

$$= (1-W_x)(1-W_y)\left(1+\frac{\varepsilon\delta\lambda_{WT}}{(1-\varepsilon)\kappa_t+(1-\delta)\kappa_n}\right). \tag{15}$$

We see a recursive relationship appear, such that

$$\frac{\varepsilon\delta\lambda_{WT}}{(1-\varepsilon)\kappa_t+(1-\delta)\kappa_n} = (1-W_x)(1-W_y)\left(1+(1-W_x)(1-W_y)\left(1+...\right)\right) \tag{16}$$

$$= (1-W_x)(1-W_y)+((1-W_x)(1-W_y))^2+((1-W_x)(1-W_y))^3+... \tag{17}$$

Therefore, we have from Eq. (8) that

$$W_{xy} = W_x W_y \sum_{k=0}^{\infty}((1-W_x)(1-W_y))^k \tag{18}$$

$$= W_x W_y \frac{1}{1-(1-W_x)(1-W_y)} \tag{19}$$

$$= W_x W_y \left(1+\frac{(1-W_x)(1-W_y)}{1-(1-W_x)(1-W_y)}\right), \tag{20}$$

from the property of geometric series and noting that $(1-W_x)(1-W_y) \leq 1$.

##### 1.2.2 Scott-Hwa model with no feedback

To understand the impact of growth rate optimization in the Scott-Hwa model, we consider an alternative model formulation where this feedback is absent. We assume that we still have two different processes (metabolism and translation) and that the flux through both of them has to be equal. However, we do not incorporate the assumption that the proteome has to be partitioned between these two processes. Instead, each process is assumed to have a given amount of proteins or enzymes available at its disposal. This formulation implies that

$$\lambda = \kappa_t \phi_t = \kappa_n \phi_n, \tag{21}$$

$$0 \leq \phi_t \leq \phi_t^{max}, \tag{22}$$

$$0 \leq \phi_n \leq \phi_n^{max}, \tag{23}$$

where $\kappa_t$ and $\kappa_n$ are the capacities of the translation and metabolic sector, respectively, as in the Scott-Hwa model, and where $\phi_t$ and $\phi_n$ are the normalized concentrations of proteins or enzymes.

If we assume that the cell maximizes its growth rate, we have that

$$\lambda_{max}^{nf} = \min(\kappa_t \phi_t^{max}, \kappa_n \phi_n^{max}), \tag{24}$$

where the $nf$ superscript stands for *no feedback*.

Clearly, in this simplified model, one sector will be limiting either because it is not efficient enough, i.e., $\kappa_i$ is too small, or because it does not have enough resources to deploy to accelerate the process, i.e., $\phi_i^{max}$ is too small. In this case, we see that the growth rate $\lambda$ corresponds to the minimum of the potential maximal fluxes in either sector.

Therefore, modeling mutations as affecting either $\kappa_t$ or $\kappa_n$ but not the protein fractions, we find that

$$W_x = \frac{\min(\kappa_t' \phi_t^{max}, \kappa_n \phi_n^{max})}{\lambda_0}$$

$$W_y = \frac{\min(\kappa_t \phi_t^{max}, \kappa_n' \phi_n^{max})}{\lambda_0}$$

$$W_{xy} = \frac{\min(\kappa_t' \phi_t^{max}, \kappa_n' \phi_n^{max})}{\lambda_0}$$

For the sake of argument, assume that $\kappa_t \phi_t^{max} \leq \kappa_n \phi_n^{max}$. Therefore, $\lambda_0 = \kappa_t \phi_t^{max}$ and $W_x = \frac{\kappa_t' \phi_t^{max}}{\lambda_0} = \frac{\kappa_t'}{\kappa_t}$, and $W_y = \min(1, \frac{\kappa_n' \phi_n^{max}}{\kappa_t \phi_t^{max}})$.

We can then write that

$$W_{xy} = \min\left(W_x, \frac{\kappa_n' \phi_n^{max}}{\kappa_t \phi_t^{max}}\right).$$

If $\frac{\kappa_n' \phi_n^{max}}{\kappa_t \phi_t^{max}} > 1$, then $W_{xy} = W_x \leq 1$. However, if $\frac{\kappa_n' \phi_n^{max}}{\kappa_t \phi_t^{max}} \leq 1$, then $W_y = \frac{\kappa_n' \phi_n^{max}}{\kappa_t \phi_t^{max}}$. Therefore, we have that

$$W_{xy} = \min\left(W_x, W_y\right),$$

that is, in the model with no feedback laid out above, double-mutant fitnesses are actually the minimum of the single mutant fitnesses. Simulation results are reported in Fig. S4.

We note that this absence of feedback results in significantly smaller growth rates. In the Scott-Hwa model, we have

$$\lambda^{SH} = \frac{\kappa_t \kappa_n}{\kappa_t + \kappa_n}.$$

This implies that $\phi_t = \frac{\kappa_n}{\kappa_t+\kappa_n}$ and $\phi_n = \frac{\kappa_t}{\kappa_t+\kappa_n}$.

Let us now consider, in the context of the model in this section, $\phi_t^{max} = \frac{\kappa_n}{\kappa_t+\kappa_n}$ and $\phi_n^{max} = \frac{\kappa_t}{\kappa_t+\kappa_n}$. Upon mutation, of $\kappa_t$ or $\kappa_n$, we have

$$\frac{\lambda^{nf}}{\lambda^{SH}} = \frac{\min\left(\kappa_t' \frac{\kappa_n}{\kappa_t+\kappa_n}, \kappa_n' \frac{\kappa_t}{\kappa_t+\kappa_n}\right)}{\frac{\kappa_t'\kappa_n'}{\kappa_t'+\kappa_n'}} \tag{25}$$

$$= \min\left(\frac{\kappa_n(\kappa_t'+\kappa_n')}{\kappa_n'(\kappa_t+\kappa_n)}, \frac{\kappa_t(\kappa_t'+\kappa_n')}{\kappa_t'(\kappa_t+\kappa_n)}\right) \tag{26}$$

$$= \frac{\kappa_t'+\kappa_n'}{\kappa_t+\kappa_n} \min\left(\frac{\kappa_n}{\kappa_n'}, \frac{\kappa_t}{\kappa_t'}\right) \tag{27}$$

Assume, without loss of generality, that $\frac{\kappa_t}{\kappa_t'} \leq \frac{\kappa_n}{\kappa_n'}$, i.e. $\frac{\kappa_n'}{\kappa_n} \leq \frac{\kappa_t'}{\kappa_t}$. Then, we have

$$\frac{\kappa_t'+\kappa_n'}{\kappa_t+\kappa_n} \leq \frac{\kappa_t'+\kappa_t'\frac{\kappa_n}{\kappa_t}}{\kappa_t+\kappa_n} = \frac{\kappa_t'}{\kappa_t},$$

which implies that $\lambda^{nf}/\lambda^{SH} \leq 1$. Therefore, in the absence of feedback, the growth rate is smaller than in the Scott-Hwa model, which is consistent with the interpretation of growth-rate optimization.

#### 1.3 Weiße model

##### 1.3.1 Original model

The original model consists of a system of equations describing the synthesis, degradation, and reaction of a series of molecules and proteins. In particular, external nutrients $s$ are converted into internal nutrients $s_i$. Those nutrients are then converted into a generic cellular energy $a$ that enables transcription and translation. Indeed, mRNAs $m$ are transcribed and then bind to ribosomes to form an mRNA-ribosome complex $c$ that governs the rate of protein translation. There are four main types of mRNAs and complexes in the model, indicated by a subscript. Each are associated to different processes: transport, metabolism, ribosomal, and q-proteins. This last type of protein denotes housekeeping proteins whose concentration remains approximately constant.

$$\dot{s_i} = \nu_{\text{imp}}(e_t, s) - \nu_{\text{cat}}(e_m, s_i) - \lambda s_i \tag{28}$$

$$\dot{a} = n_s \cdot \nu_{\text{cat}}(e_m, s_i) - \sum_{x\in\{r,t,m,q\}} n_x \nu_x(c_x, a) - \lambda a \tag{29}$$

$$\dot{r} = \nu_r(c_r, a) - \lambda r + \sum_{x\in\{r,t,m,q\}} (\nu_x(c_x, a) - k_b r m_x + k_u c_x) \tag{30}$$

$$\dot{e_t} = \nu_t(c_t, a) - \lambda e_t \tag{31}$$

$$\dot{e_m} = \nu_m(c_m, a) - \lambda e_m \tag{32}$$

$$\dot{q} = \nu_q(c_q, a) - \lambda q \tag{33}$$

$$\dot{m_x} = \omega_x(a) - (\lambda + d_m)m_x + \nu_x(c_x, a) - k_b r m_x + k_u c_x \tag{34}$$

$$\dot{c_x} = -\lambda c_x + k_b r m_x - k_u c_x - \nu_x(c_x, a) \tag{35}$$

All of these reactions are modulated by parameters reported in Table S2.

| | description | default value | unit |
|---|---|---|---|
| $s$ | external nutrient | $10^4$ | [molecs] |
| $d_m$ | mRNA-degradation rate | 0.1 | [$\text{min}^{-1}$] |
| $n_s$ | nutrient efficiency | 0.5 | none |
| $n_r$ | ribosome length | 7459 | [aa/molecs] |
| $n_x, x \in \{t, m, q\}$ | length of non-ribosomal proteins | 300 | [aa/molecs] |
| $\gamma_{\max}$ | max. transl. elongation rate | 1260 | [aa/min molecs] |
| $K_\gamma$ | transl. elongation threshold | 7 | [molecs/cell] |
| $v_t$ | max. nutrient import rate | 726 | [$\text{min}^{-1}$] |
| $K_t$ | nutrient import threshold | 1000 | [molecs] |
| $v_m$ | max. enzymatic rate | 5800 | [$\text{min}^{-1}$] |
| $K_m$ | enzymatic threshold | 1000 | [molecs/cell] |
| $w_r$ | max. ribosome transcription rate | 930 | [molecs/min cell] |
| $w_e = w_t = w_m$ | max. enzyme transcription rate | 4.14 | [molecs/min cell] |
| $w_q$ | max. q-transcription rate | 948.93 | [molecs/min cell] |
| $\theta_r$ | ribosome transcription threshold | 426.87 | [molecs/cell] |
| $\theta_{nr}$ | non-ribosomal transcription threshold | 4.38 | [molecs/cell] |
| $K_q$ | q-autoinhibition threshold | 152 219 | [molecs/cell] |
| $h_q$ | q-autoinhibition Hill coeff. | 4 | none |
| $k_b$ | mRNA-ribosome binding rate | 1 | [cell/min molecs] |
| $k_u$ | mRNA-ribosome unbinding rate | 1 | [$\text{min}^{-1}$] |
| $M$ | total cell mass | $10^8$ | [aa] |

Table S2: Model parameters from (Weiße et al. 2015), obtained either from the literature or from parameter optimization.

The rates governing the system of equations are assumed to follow Michaelis-Menten kinetics as follows

$$\nu_{\text{imp}}(e_t, s) = e_t \frac{v_t s}{K_t + s} \tag{36}$$

$$\nu_{\text{cat}}(e_m, s_i) = e_m \frac{v_m s_i}{K_m + s_i} \tag{37}$$

$$\nu_x(c_x, a) \sim c_x \frac{\gamma(a)}{n_x} \tag{38}$$

$$\gamma(a) := \frac{\gamma_{max} a}{K_\gamma + a} \tag{39}$$

$$\omega_x(a) = \omega_x \frac{a}{\theta_x + a} \tag{40}$$

Finally, the growth rate corresponds to total mass of proteins being synthesized at steady-state, that is

$$\lambda = \frac{\gamma(a) \sum_x c_x}{M}. \tag{41}$$

##### 1.3.2 Isolation of parameters

This model has a total of 21 parameters whose values were set (either from the literature or estimated) in the original model (Table S2 in (Weiße et al. 2015), reproduced here under Table S2).

Among them, some represent quantities or parameters that would not reasonably change under a mutation. Therefore, we do not consider the following parameters in our mutation analysis: external nutrients $s$, mRNA degradation rate $d_m$, ribosome length $n_r$, length of non-ribosomal proteins $n_x$, q-autoinhibition Hill coefficient $h_q$, the mRNA-ribosome binding/unbinding rates $k_b$ and $k_u$, the total cell mass $M$.

To determine whether a numerical mutational analysis with the remaining 13 parameters could be considered, we computed the impact of a change of these parameters on the growth rate $\lambda$ (Fig. S6). Upon mutation, most parameters do impact the growth rate negatively. However, two parameters $v_m, K_m$ do not seem to have an impact on the growth rate. We attribute this to the metabolic sector not being limiting in this model, with those parameter values. We therefore exclude those two parameters from further analysis. We also exclude the parameters associated to the q-proteins $K_q$ and $\omega_q$ as these are associated mostly to housekeeping functions.

We are therefore left with 9 parameters modeling the impact of 4 different processes. In total, this results in 28 potential combinations of two parameters associated with different processes.

##### 1.3.3 Simplification

To facilitate analytical treatment, we consider a simplified model with only two types of proteins, instead of four: a general protein $p$ and ribosomes $r$. We also eliminate the equations corresponding to the competitive binding of mRNAs for ribosomes. Practically, this assumption means that every mRNA binds immediately to a ribosome. While this assumption might not be true in all regimes, we find that this simplifies the analytical treatment and still enables us to understand something concrete about the model.

The simplified model we consider is

$$\dot{s}_i = \nu_{\text{imp}}(p, s) - \nu_{\text{cat}}(p, s_i) - \lambda s_i \tag{42}$$

$$\dot{a} = n_s \nu_{\text{cat}}(p, s_i) - \sum_x n_x \nu_x(c_p, a) - \lambda a \tag{43}$$

$$\dot{r} = \nu_r(c_r, a) - \lambda r + \sum_x \nu_x(c_x, a) \tag{44}$$

$$\dot{p} = \nu_p(p, a) - \lambda p \tag{45}$$

$$\dot{c}_r = \omega_r(a) - \nu_r(c_r, a) - \lambda c_r \tag{46}$$

$$\dot{c}_p = \omega_p(a) - \nu_p(c_p, a) - \lambda c_p \tag{47}$$

If we assume the dilution fluxes to be negligible in comparison to other fluxes in Eqs..., we get that

$$\nu_{\text{imp}}(p, s) \sim \nu_{\text{cat}}(p, s_i) \Rightarrow p v_t \frac{s}{K_t + s} \sim p v_m \frac{s_i}{K_m + s_i} \tag{48}$$

$$n_s \nu_{\text{cat}}(p, s_i) \sim \sum_x n_x \nu_x(c_p, a) = (c_p + c_r)\gamma(a) \tag{49}$$

$$\omega_p(a) \sim \nu_p(c_p, a) \tag{50}$$

$$\omega_r(a) \sim \nu_r(c_r, a) \tag{51}$$

The dilution flux needs to be nonzero for $p$, as its governing equation only contains two terms. We have therefore that

$$\nu_p(p, a) = \lambda p \Rightarrow p \sim \omega_p \frac{a}{\theta_x + a} \frac{1}{\lambda} \tag{52}$$

As the growth rate $\lambda = \frac{\gamma(a) \sum_x c_x}{M}$, we have

$$\lambda = \frac{n_s \nu_{\text{cat}}}{M} \sim \frac{1}{M} n_s v_t p \frac{s}{K_t + s} \tag{53}$$

Injecting Eq. (52) into (53), we have

$$\lambda \sim \sqrt{\frac{1}{M} n_s v_t w_p \frac{s}{K_t + s} \frac{a}{\theta_x + a}} \tag{54}$$

From the above expression, we see that terms belonging to different sectors combine as a product. However, the energy $a$ is **not** a parameter, but rather a variable in the model. Its value also directly depends on the values of other parameters.

From Eq. (49), we have that

$$\lambda \sim \frac{1}{M} \left( n_p \omega_p \frac{a}{\theta_x + a} + n_r \omega_r \frac{a}{\theta_r + a} \right) \tag{55}$$

If $\theta_x < a < \theta_r$, we can approximate that with

$$\lambda \sim \frac{1}{M} \left( n_p \omega_p + n_r \omega_r \frac{a}{\theta_r} \right). \tag{56}$$

If $a \ll \theta_x, \theta_r$, we have

$$\lambda \sim \frac{1}{M} \left( \frac{n_p \omega_p}{\theta_x} + \frac{n_r \omega_r}{\theta_r} \right) a. \tag{57}$$

In both cases, we can assume that there is a linear relationship between the growth rate $\lambda$ and the energy vector $a$, such that

$$a \sim b\lambda + c. \tag{58}$$

We can inject that in the $a/(\theta_x + a)$ term in Eq. (54), which gives

$$\frac{a}{\theta_x + a} \sim \frac{\lambda + c/b}{(\theta_x + c)/b + \lambda} \tag{59}$$

Assuming that $c/b \ll \lambda$, we have

$$\frac{a}{\theta_x + a} \sim \frac{\lambda}{K' + \lambda}, \tag{60}$$

with $K' = (\theta_x + c)/b$.

Injecting the above in Eq. (54), we get that,

$$\lambda \sim \sqrt{\frac{1}{M} n_s v_t w_p \frac{s}{K_t + s} \frac{\lambda}{K' + \lambda}}. \tag{61}$$

Therefore, $K'$ acts as a scale of growth rate where deviations from the product model can appear. If $K' \gg \lambda$, then

$$\lambda \sim \frac{1}{M} n_s v_t w_p \frac{s}{K_t + s}. \tag{62}$$

On the other hand, if $K' \ll \lambda$, then

$$\lambda \sim \sqrt{\frac{1}{M} n_s v_t w_p \frac{s}{K_t + s}}. \tag{63}$$

In both of these cases, the growth rate $\lambda$ behaves as the product of multiple parameters. However, if $\lambda \sim K'$, and if we denote two parameters in the expression by $\alpha, \beta$ without loss of generality, we have

$$\lambda^2 (K' + \lambda) \sim \alpha\beta\lambda. \tag{64}$$

Either $\lambda = 0$ (which we will not consider), or

$$\begin{aligned} \lambda(K' + \lambda) &= \alpha\beta \\ \Leftrightarrow \lambda^2 + K'\lambda - \alpha\beta &= 0 \\ \Leftrightarrow \lambda &= \frac{-K' + \sqrt{K'^2 + 4\alpha\beta}}{2} \\ &= \frac{K'}{2}\left(-1 + \sqrt{1 + \frac{4\alpha\beta}{K'^2}}\right) \end{aligned}$$

We can write, for small $x$, that

$$\sqrt{1+x} = 1 + \frac{x}{2} - \frac{x^2}{8} + \dots$$

Assuming that $\frac{4\alpha\beta}{K'^2}$ is small and keeping terms up to second order, we therefore have that

$$\begin{aligned} \lambda &= \frac{K'}{2}\frac{x}{2}\left(1 - \frac{x}{4}\right), \text{ with } x = \frac{4\alpha\beta}{K'^2} \\ \Rightarrow \lambda &\propto \alpha\beta\left(1 - \gamma\alpha\beta\right), \text{ with } \gamma = 1/K'^2 \end{aligned}$$

##### 1.4 Double mutant fitness

From there, we can compute an approximation to the double mutant fitness. Let us assume that $\lambda = \alpha\beta\left(1 - \gamma\alpha\beta\right)$ as derived above. To simplify notation, we will rescale the parameters to their $WT$ value, i.e.

$$\begin{aligned} \alpha' &= \alpha/\alpha_{WT}, \\ \beta' &= \beta/\beta_{WT}, \\ \bar{\gamma} &= \gamma\alpha_{WT}\beta_{WT}. \end{aligned}$$

Under this rescaling, we see that

$$\begin{aligned} \lambda_{WT} &= \alpha_{WT}\beta_{WT}(1 - \bar{\gamma}), \\ W_x &= \frac{\alpha'(1 - \bar{\gamma}\alpha')}{1 - \bar{\gamma}}, \\ W_y &= \frac{\beta'(1 - \bar{\gamma}\beta')}{1 - \bar{\gamma}}, \\ W_{xy} &= \alpha'\beta'\frac{1 - \bar{\gamma}\alpha'\beta'}{(1 - \bar{\gamma})}. \end{aligned}$$

Noting that

$$W_x W_y = \frac{\alpha'\beta'(1-\bar{\gamma}\alpha')(1-\bar{\gamma}\beta')}{(1-\bar{\gamma})^2}$$

we can show that

$$\begin{aligned} W_{xy} &= W_x W_y - \frac{\alpha'\beta'}{(1-\bar{\gamma})^2}\bar{\gamma}(1-\alpha')(1-\beta'), \\ &= W_x W_y \left(1 - \bar{\gamma}\frac{1-\alpha'}{1-\bar{\gamma}\alpha'}\frac{1-\beta'}{1-\bar{\gamma}\beta'}\right). \end{aligned}$$

If $\bar{\gamma}$ is small, then $W_x = \frac{\alpha'(1-\bar{\gamma}\alpha')}{1-\bar{\gamma}} \sim \alpha'$ and $W_y = \frac{\beta'(1-\bar{\gamma}\beta')}{1-\bar{\gamma}} \sim \beta'$, and therefore we can write that

$$W_{xy} \sim W_x W_y \left(1 - \bar{\gamma}\frac{1-W_x}{1-\bar{\gamma}W_x}\frac{1-W_y}{1-\bar{\gamma}W_y}\right).$$

### 2 Supplementary Figures

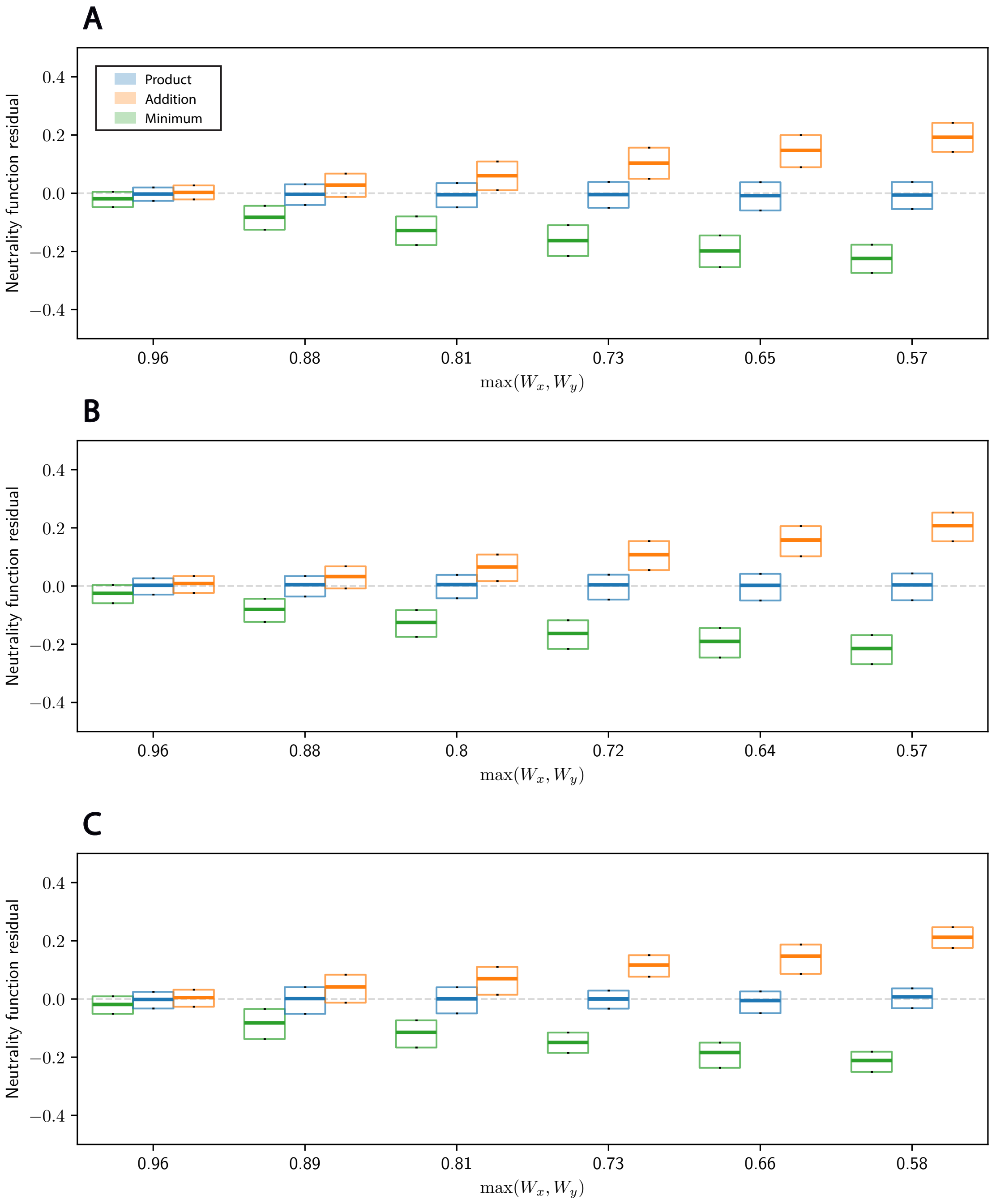


Figure S1: **Double mutant fitnesses are best described by the Product neutrality function in the SGA dataset.** Box plots for the distributions of the residuals for the three neutrality functions as a function of the maximum single mutant fitness. Each plot corresponds to a different subset of the SGA dataset. Namely, they correspond to the first set of query mutants (see Methods) crossed to different types of mutant arrays in different temperature conditions (**A.** Deletion Mutant Array at 26°C. **B.** Temperature Sensitive Array at 26°C. **C.** Temperature Sensitive Array at 30°C.). Thick line denotes the median, and boxes denote the 25th and 75th percentiles of the distributions. The Product neutrality function models the data consistently better than the others.

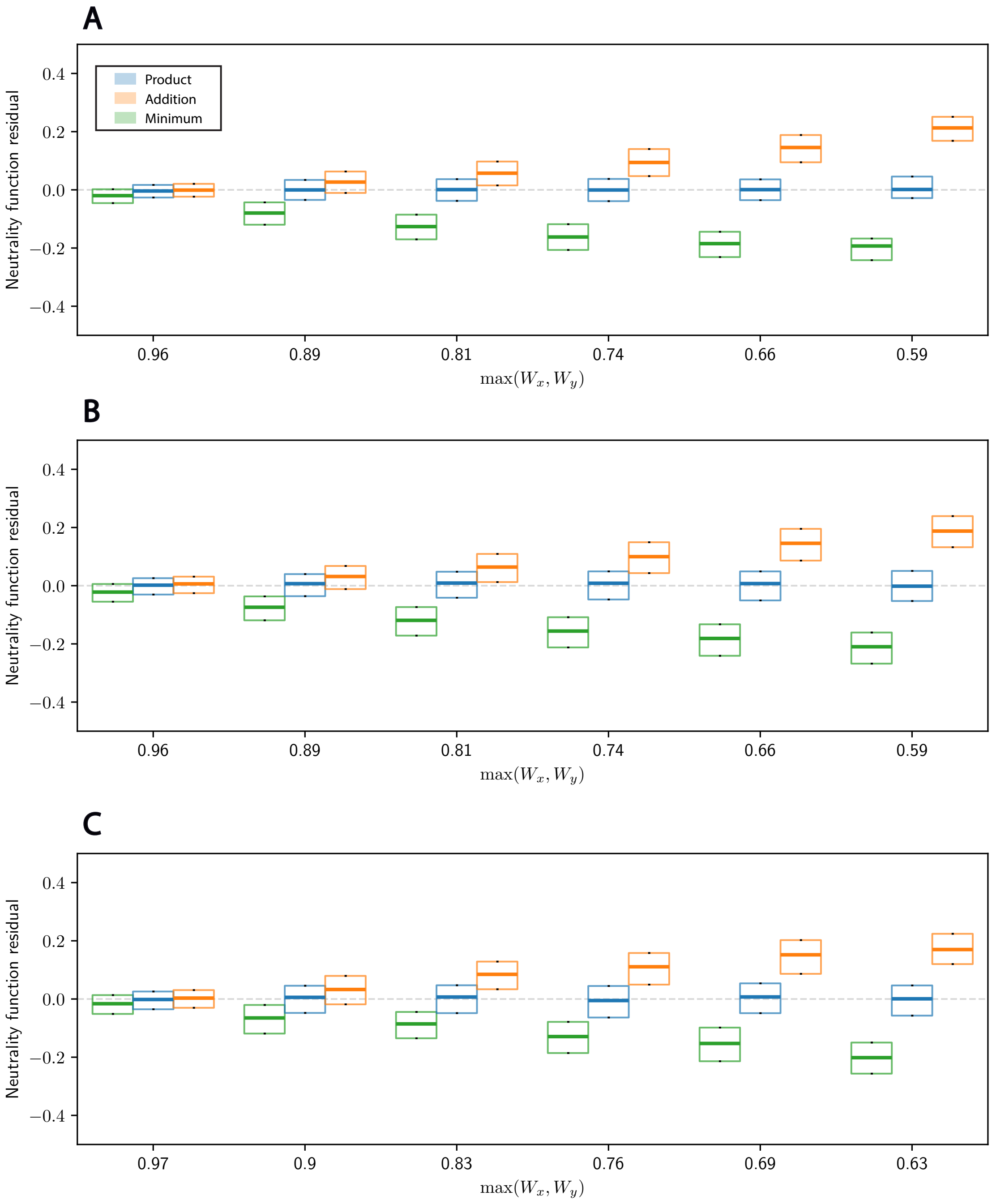


Figure S2: **Double mutant fitnesses are best described by the Product neutrality function in the SGA dataset.** Box plots for the distributions of the residuals for the three neutrality functions as a function of the maximum single mutant fitness. Each plot corresponds to a different subset of the SGA dataset. Namely, they correspond to the second set of query mutants (DAmP, see Methods) crossed to different types of mutant arrays in different temperature conditions (**A.** Deletion Mutant Array at 30°C. **B.** Temperature Sensitive Array at 26°C. **C.** Temperature Sensitive Array at 30°C.). Thick line denotes the median, and boxes denote the 25th and 75th percentiles of the distributions. The Product neutrality function models the data consistently better than the others.

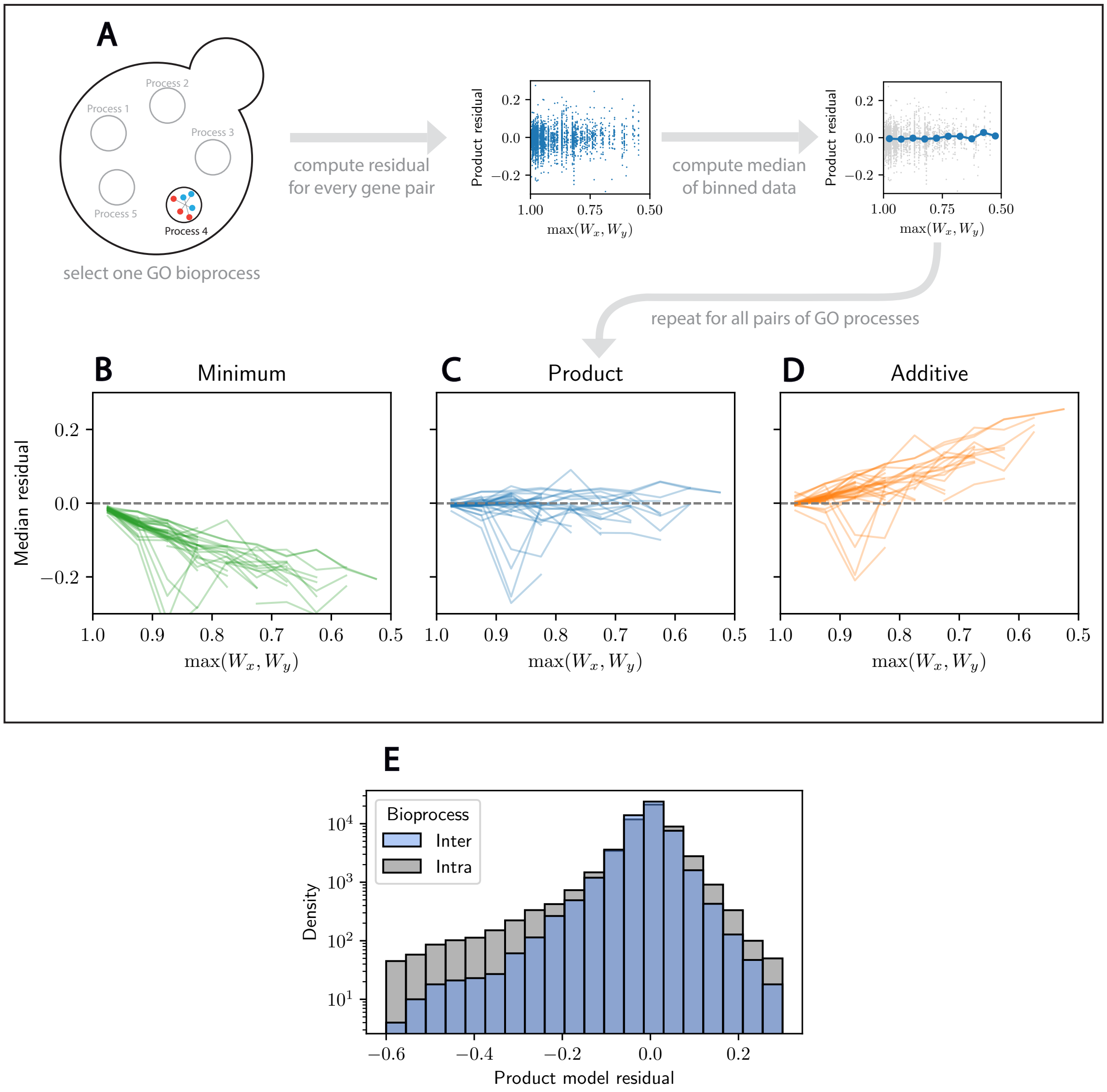


Figure S3: **Larger deviations from the Product neutrality function characterize gene pairs affecting the same GO biological process.** **A.** Schematic illustration of the analysis process for double mutants where both mutations affect the same GO biological process. We first select two different GO biological processes and extract the double mutants in the SGA dataset associated with them. Then, we compute the median residual for each pair of biological processes and each neutrality function. **B-D.** Median residual for the Minimum, Product, and Additive neutrality functions as a function of the maximum single mutant fitness. Each line denotes mutations to a single GO biological process. We see larger deviations from the Product model than in Fig. 2. **E.** Histogram of the SGA dataset after extracting pairs affecting either two different (inter) or the same (intra) GO biological process. Large residuals are much more likely when both mutations affect the same GO biological process.

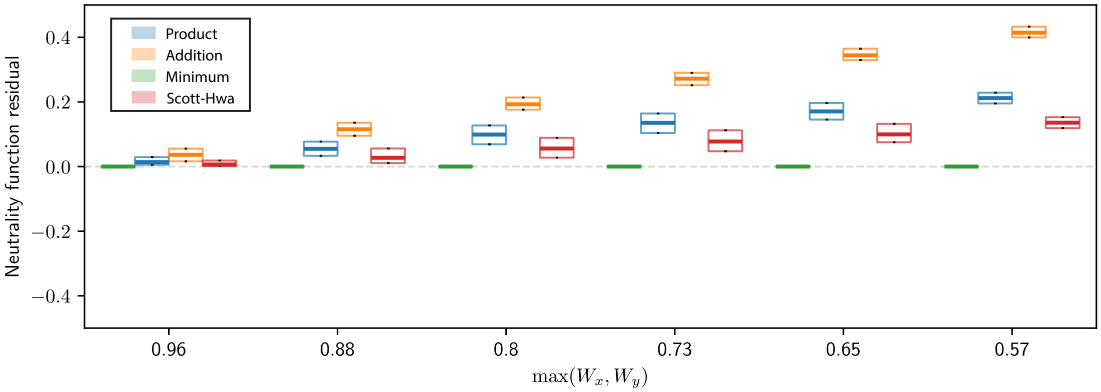


Figure S4: **The Scott-Hwa model with no feedback follows a Minimum neutrality function.** Box plots for the distributions of the residuals for the model of Section 1.2.2 as a function of the maximum single mutant fitness. Thick line denotes the median, and boxes denote the upper and lower quartiles of the data. The absence of feedback due to resource competition in the model of Section 1.2.2 results in a Minimum neutrality function.

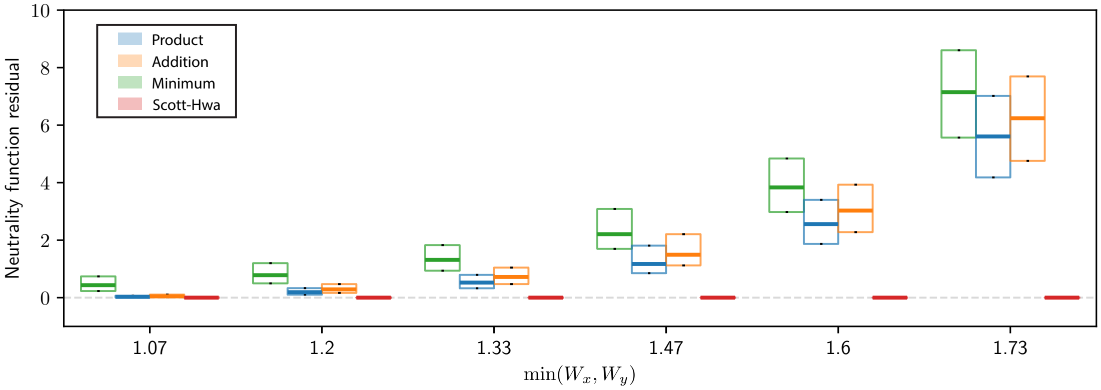


Figure S5: **Large deviations from the Product neutrality function characterize beneficial mutations.** Box plots for the distributions of the residuals for the different neutrality functions for beneficial mutations as a function of the minimum single mutant fitness. Thick lines denote the median, and boxes denote the upper and lower quartiles of the data. Deviations from the Product neutrality function derived in Section 1.2.1 can be unbounded.

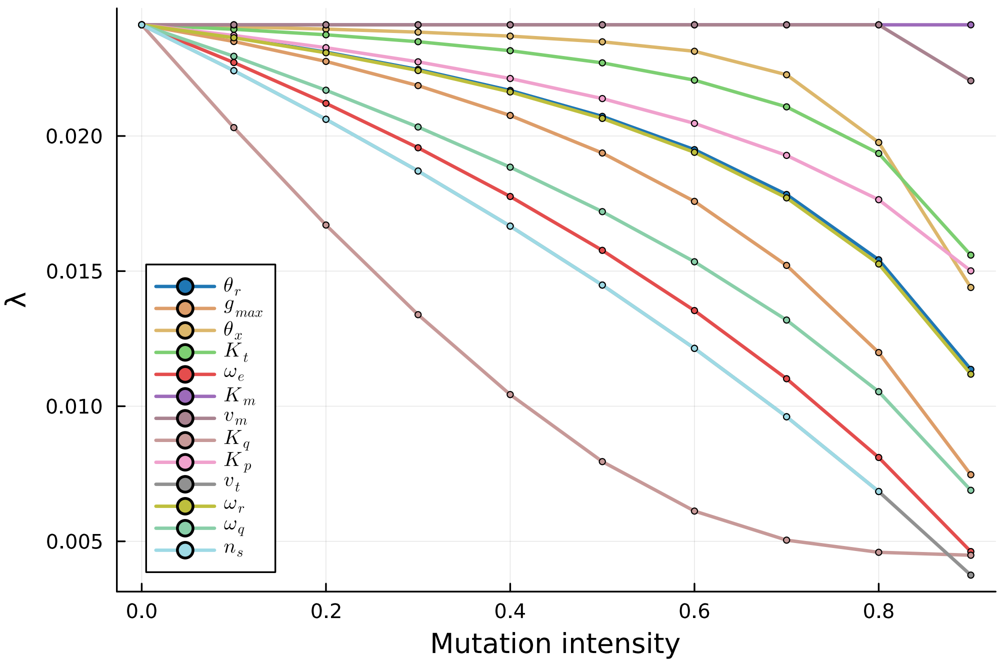


Figure S6: **11 parameters exhibit negative impact on growth rate upon mutation in the Weiße model.** Among the initial 21 parameters, 13 were kept as candidates for a mutational analysis. Two of them $v_m, K_m$, associated with the metabolic sector, do not have any impact on growth rate upon mutation, likely because this sector is not limiting for growth in that parameter range. The others have a negative impact upon mutation.

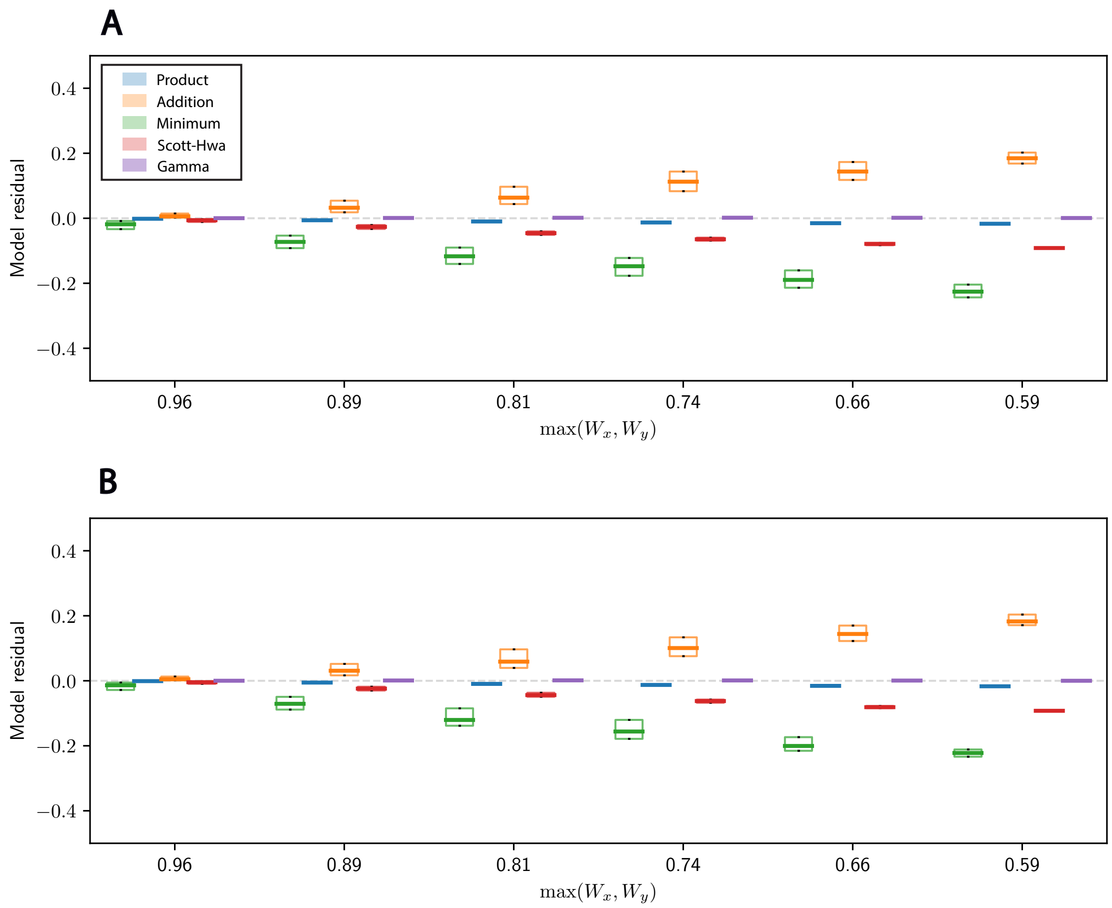


Figure S7: **Deviations from the Product neutrality function in the Weiße model are captured by the $\gamma$ approximation.** Box plots for the distributions of the residuals for all models considered in this paper, for two example parameter pairs (**A.** $\omega_q, n_s$, **B.** $K_t, \omega_e$.). Thick line denotes the median, and boxes denote the upper and lower quartiles of the data. The Gamma model, in purple, denotes the derivation in Fig. 5A. It captures the small deviations from the Product neutrality function.

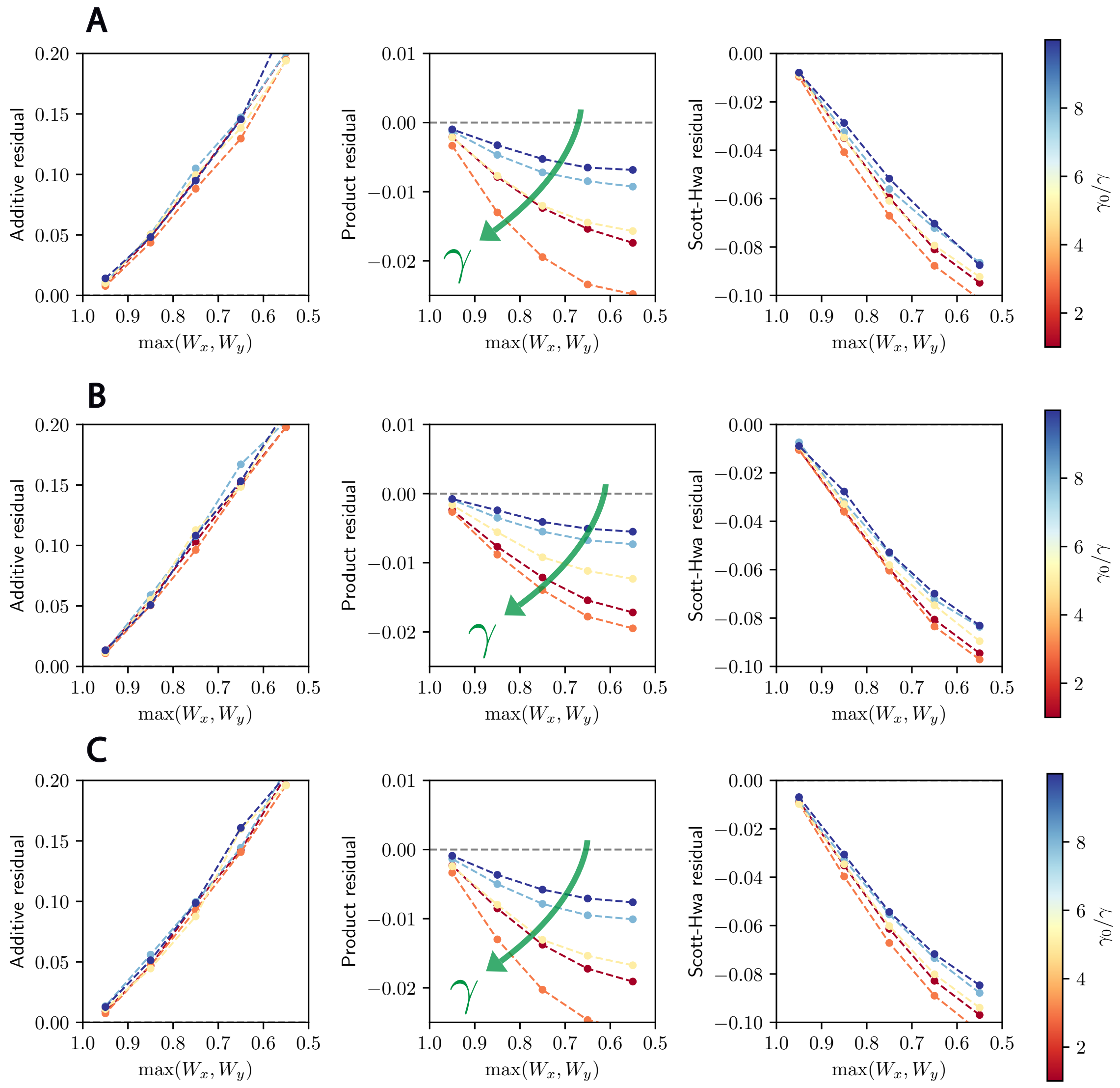


Figure S8: **Tuning $\gamma$ impacts how good an approximation the Product neutrality function is for multiple parameter pairs in the Weiße model.** The analysis of Fig. 5C, illustrating the impact of $\gamma$ on a single parameter pair $(n_s, v_t)$ is here extended to other parameter pairs to demonstrate the validity of the mechanistic interpretation. **A.** $v_t, \omega_r$; **B.** $\omega_e, n_s$; **C.** $\theta_r, v_t$. In all cases, we see that decreasing $\gamma$ results in better alignment with the Product model.
